# OncoLoop: A network-based precision cancer medicine framework

**DOI:** 10.1101/2022.02.11.479456

**Authors:** Alessandro Vasciaveo, Min Zou, Juan Martín Arriaga, Francisca Nunes de Almeida, Eugene F. Douglass, Maho Shibata, Antonio Rodriguez-Calero, Simone de Brot, Antonina Mitrofanova, Chee Wai Chua, Charles Karan, Ron Realubit, Sergey Pampou, Jaime Y. Kim, Eva Corey, Mariano J. Alvarez, Mark A. Rubin, Michael M. Shen, Andrea Califano, Cory Abate-Shen

## Abstract

At present, prioritizing cancer treatments at the individual patient level remains challenging, and performing co-clinical studies using patient-derived models in real-time is often not feasible. To circumvent these challenges, we introduce OncoLoop, a precision medicine framework to predict and validate drug sensitivity in human tumors and their pre-existing high-fidelity (*cognate*) model(s) by leveraging perturbational profiles of clinically-relevant oncology drugs. As proof-of-concept, we applied OncoLoop to prostate cancer (PCa) using a series of genetically-engineered mouse models (GEMMs) that recapitulate a broad spectrum of disease states, including castration-resistant, metastatic, and neuroendocrine prostate cancer. Interrogation of published cohorts using Master Regulator (MR) conservation analysis revealed that most patients were represented by at least one cognate GEMM-derived tumor (GEMM-DT). Drugs predicted to invert MR activity in patients and their cognate GEMM-DTs were successfully validated, including in two cognate allografts and one cognate patient-derived xenograft (PDX). OncoLoop is a highly generalizable framework that can be extended to other cancers and potentially other diseases.

**Significance Statement:** OncoLoop is a transcriptomic-based experimental and computational framework that can support rapid-turnaround co-clinical studies to identify and validate drugs for individual patients, which can then be readily adapted to clinical practice. This framework should be applicable in many cancer contexts for which appropriate models and drug perturbation data are available.

## Introduction

Systematic prediction of drug efficacy *in vivo* remains a major clinical challenge for most cancer types. This is due, in part, to tumor heterogeneity, which makes it difficult to optimize treatments on an individual basis. This can be further compounded by difficulties in establishing patient-derived models that recapitulate the biology and complexity of an individual patient’s tumor for co-clinical validation. Indeed, for some tumor types, establishment of patient-derived xenograft (PDX) models can take several months to more than one year (1, 2), thus compromising the ability to use these models to predict and validate drug efficacy within a time-frame that is compatible with patient treatment, especially in the aggressive metastatic setting. Patient-derived organoid models have become increasingly more accessible and representative; however, these may be limited in their ability to model the tumor microenvironment (3, 4). Finally, although human cell lines derived from patient’s tumors or metastases are widely available for many cancers, they rarely represent the full spectrum of cancer phenotypes observed in patients and often have idiosyncratic dependencies, as a result of genetic and epigenetic alterations they may accrue to survive *in vitro*. In principle, genetically-engineered mouse models (GEMMs), which are now widely available for many cancer types (5), can provide valuable resources for studying drug response in the whole organism in the context of the native tumor microenvironment. However, their effective use in co-clinical studies has been limited, since few studies have assessed their fidelity to their human counterparts, in terms of recapitulating the biology and drug sensitivity of patient tumors (6).

These general challenges are exemplified in prostate cancer (PCa), which remains the most prevalent form of cancer and a leading cause of cancer-related death in men (7). PCa is characterized by its broad range of disease outcomes; in particular, while men with locally-invasive disease, which account for the vast majority of new diagnoses, have a 5-year survival of >90%, those who progress to advanced PCa have a 5-year survival of < 30%. The first line treatment for advanced PCa is androgen deprivation therapy (ADT), which initially leads to tumor regression but ultimately to the emergence of castration-resistant prostate cancer (CRPC), which is often metastatic (mCRPC) (8–10). Second line treatments include taxane-based chemotherapy and anti-androgens, such as Enzalutamide and Abiraterone; while these may initially be effective, many patients fail treatment, which is associated with the emergence of highly aggressive disease variants, including neuroendocrine prostate cancer (NEPC) (11, 12). Challenges in treating advanced PCa include its inherent molecular heterogeneity, the infrequency of driver mutations, and its long oncogenic tail (13), which make it difficult to determine *a priori* which specific treatments are likely to effective for a given patient. Furthermore, while there are many GEMMs for prostate cancer (14), it is proven difficult to model human PCa since the establishment of patient-derived xenograft (PDX) as well as patient-derived organoid models has been particularly low yield, especially for advanced disease stages (15–19).

To overcome these challenges, we developed OncoLoop, a framework that integrates experimental and computational data to first identify high-fidelity models for a given human tumor (*cognate* model, hereafter), and then to predict optimal drug treatments for the patient and for its cognate model based on large-scale drug perturbation profiles, and lastly to validate drug efficacy using the cognate model. The rationale for OncoLoop is based our previous studies in which we have shown that MR proteins represent mechanistic determinants of a tumor’s transcriptional state, while inversion of their activity effectively abrogates tumor viability (20–22). Thus, cognate models for a given patient are identified by assessing Master Regulator (MR) protein activity conservation for a patient and cognate model pair (23, 24); conversely, optimal drugs are prioritized based on their ability to invert the MR-activity signature of both the patient tumor and its cognate model (21). Hence, OncoLoop identifies the most statistically significant MR three-way relationships encompassing MR-matched tumors and their highest-fidelity cognate model(s), and one or more drugs predicted to invert their MR-activity signature (hereafter, MR-inverter drugs).

We established OncoLoop in the context of PCa, for which both large-scale human patient cohorts—comprising both primary tumors and metastases (25, 26)—and an extensive repertoire of GEMMs ((14) and this report) are available. These analyses revealed that a majority of patients in these cohorts had at least one cognate GEMM-derived tumor (GEMM-DT). To identify MR-inverter drugs, we leveraged PanACEA (27), a large collection of drug perturbational profiles in cell lines matched to 23 tumor subtypes, including PCa. Three out of four predicted drugs prioritized by our analyses induced highly significant growth inhibition of tumor allografts from the cognate GEMM-DT *in vivo*. We further validated these predicted drugs as MR-inverters for a cognate PCa PDX model, which resulted in complete abrogation of tumor growth. Taken together our findings show that OncoLoop provides an effective framework for the rapid identification and evaluation of patient-relevant drugs in pre-existing cognate models, thus supporting its co-clinical application.

## Results

### Conceptual overview of OncoLoop

We developed OncoLoop for the purpose of identifying drugs poised to benefit patients, whose response could be evaluated in pre-existing co-clinical models (Fig. 1A). OncoLoop leverages regulatory networks—reverse-engineered from large, tumor-specific RNA-seq profile datasets—to first identify cognate models (i.e., GEMM-DTs), based on their MR-activity signature conservation with a human tumor. The second step is to prioritize candidate drugs by identifying those that invert these MR activity signatures (i.e., MR-inverter drugs) for both the human tumor and its cognate GEMM-DT, based on drug perturbation profiles of tumor-related cell lines. The third step is to validate the efficacy of the predicted drugs *in vivo* using the cognate models. We demonstrate the feasibility of OncoLoop based on PCa, for which we have generated a large series of GEMMs; however, for other cancer types PDX models may be used if large, representative collections are available.

**Figure 1:**
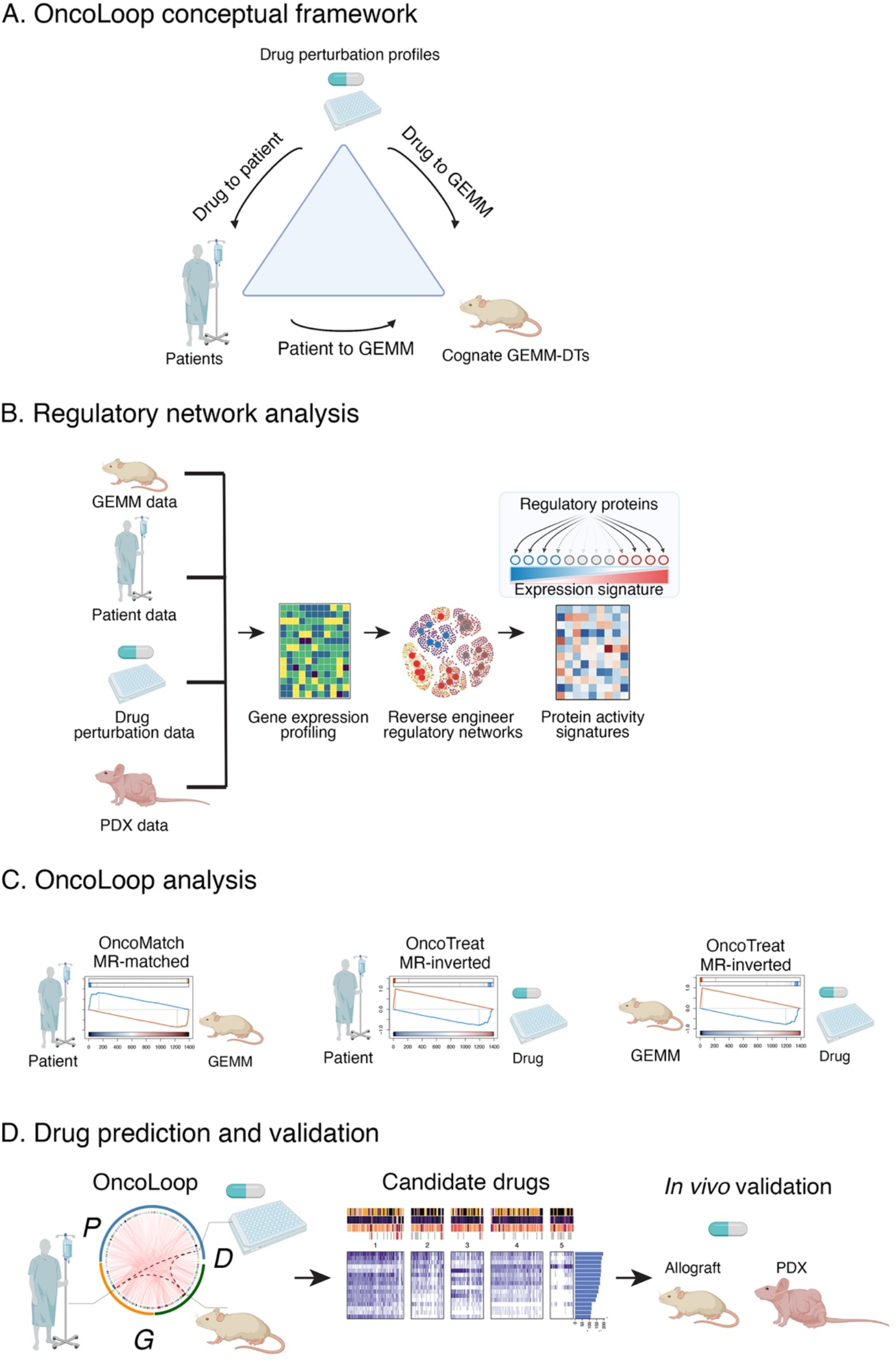
The OncoLoop conceptual framework. **A.** <u>Conceptual overview</u>: OncoLoop was designed to identify high-fidelity (cognate) models—*i.e.*, GEMM-derived tumors (GEMM-DTs), in this study— of a patient’s tumor as well as drugs capable of inverting the activity of MR proteins for both the patient and his cognate GEMM-DT. To accomplish this, OncoLoop performs integrative analysis of transcriptomic (RNA-seq) profiles from a patient’s tumor, his cognate model, and drug perturbation assays. **B.** <u>Regulatory network analysis</u>: Gene expression profiles generated from each data source are used to reverse-engineer species- and cohort-specific regulatory networks, which are then used to transform differential gene expression signature into differential protein activity profiles. **C.** <u>OncoLoop analysis</u>: Gene Set Enrichment Analysis (GSEA) is used to assess the overlap in differentially active MR proteins between a human tumor and its cognate GEMM-DTs (*OncoMatch*). Similarly, GSEA is used to identify drugs capable of inverting the MR activity (*MR-inverter drugs*) for each patient and cognate GEMM-DT(s) pair. **D.** <u>Drug prediction and validation:</u> Representative Circos plot illustrating PGD-loops generated by matching a patient (P) to a GEMM-DT (G) and by identifying the MR-inverter drug (D) for both. Candidate drugs are first prioritized by pharmacotype analysis to identify the subset of patients predicted to be sensitive to the same subset of drugs, and then validated *in vivo* using both cognate GEMM-DT-derived allografts as well as cognate PDX models.

#### Data generation

We leveraged large scale RNA-seq profiles from: (i) our comprehensive series of GEMMs representing a broad spectrum of PCa phenotypes (this study); (ii) publicly-available RNA-seq profiles from primary tumors in The Cancer Genome Atlas (TCGA) (26) and metastases in the Stand Up to Cancer-Prostate Cancer Foundation (SU2C) (25) cohorts; (iii) large-scale drug perturbation profiles from tumor-related cell lines (27); and (iv) RNA-seq profiles from well-characterized human PCa PDX models (this study) (Fig. 1B).

#### Protein activity analysis

OncoLoop requires accurate protein activity assessment for appraisal of MR-activity signatures, identification of cognate models for human tumors, and prediction of MR-inverter drugs. This is accomplished using the VIPER algorithm (23), which transforms RNA-seq profiles into accurate protein activity profiles, as recently validated by antibody-based protein abundance measurements (28). Akin to a highly multiplexed gene reporter assay, VIPER uses the expression of a protein’s tissue-specific targets (*regulon*) to measure its activity. The repertoire of targets of all regulatory and signaling proteins in a specific tissue context (*i.e.*, *context-specific interactome*) is generated by reverse-engineering large-scale, tissue-specific RNA-seq profiles using the ARACNe algorithm (29). Notably, to support accurate, model-specific protein activity measurements, we generated separate interactomes from patient-, GEMM-, and PDX-specific RNA-seq datasets (Fig. 1B).

#### GEMM cohort characterization and cognate model identification

First, we analyzed VIPER-based protein activity profiles from the GEMM-DTs to identify molecularly distinct subtypes and to demonstrate their relevance to the disease spectrum of human PCa. Next, we identified cognate GEMM-DTs, based on MR-activity signature conservation, for individual human primary PCa tumors and metastases in the TCGA and SU2C cohorts, respectively (Fig. 1C). This revealed broad coverage, in which 78% and 93% of tumors and metastases, respectively, matched to at least one cognate GEMM-DT.

#### Assessing drug mechanism of action (MoA) in MR-matched cell lines

To predict drug sensitivity, we leveraged human tumor-relevant drug perturbation profiles generated for two PCa cell lines— the androgen-dependent LNCaP and the androgen-independent DU145 cell lines—that jointly provided high-fidelity models for >80% of the tumors in the TCGA cohort, based on MR-activity signature conservation (27). To focus on the more aggressive mCRPC patients, we relied on RNA-seq profiles of DU145 cells, which were treated with a library of FDA-approved and late-stage experimental oncology drugs and vehicle control (DMSO) (*i.e.*, *drug perturbation Profiles*). Finally, the proteome-wide MoA of each drug was assessed using VIPER to measure the differential protein activity in drug-treated versus control-treated cells and was used to identify optimal MR-inverter drugs for patient-and cognate GEMM-DT pairs (Fig. 1C).

#### Closing the Loop

Having “matched” each patient tumor (*P*) to a cognate GEMM-DT (*G*) and assessed each drug (*D*) as a potential MR-inverter for both tumors, we ranked all (*P*, *G*, *D*) triplets (*PGD-loops* hereafter), based on the integration of three distinct *z-*scores, assessing the statistical significance of: (i) the similarity of a patient- and GEMM-DT MR-activity signature; (ii) the drug’s MR-activity inversion as predicted from the patient tumor; and (iii) the drug’s MR-activity inversion as predicted from the cognate GEMM-DT (Fig 1C,D).

#### In vivo validation

To prioritize drug candidates of greatest translational relevance, we focused on clinically available drugs that were most frequently nominated by the analysis of the human tumor cohorts (i.e., pharmacotyping). Among these, we evaluated the ability of the top-predicted drugs to inhibit tumor growth and recapitulate the predicted MR-activity inversion *in vivo*, based on co-clinical studies in cognate allograft and PDX models (Fig. 1D). The following sections discuss each of these steps in detail.

### A GEMM resource that models PCa progression

An essential requirement for OncoLoop is the availability of pre-existing high-fidelity models—that are accurate surrogates for their counterpart human tumors—to enable co-clinical validation of predicted drugs within a time-frame relevant to the patient’s care. Toward this end, we assembled an extensive series of GEMMs that are based on genetic and/or pathway alterations that are prevalent in human PCa and thereby recapitulate a broad spectrum of human PCa phenotypes (Fig. 2A-E, Fig. S1A-C) (30–34). A complete list of GEMMs used in this study is provided in Table S1A,B; the entire GEMM series is available from the Jackson Laboratory (see Table S1).

**Figure 2:**
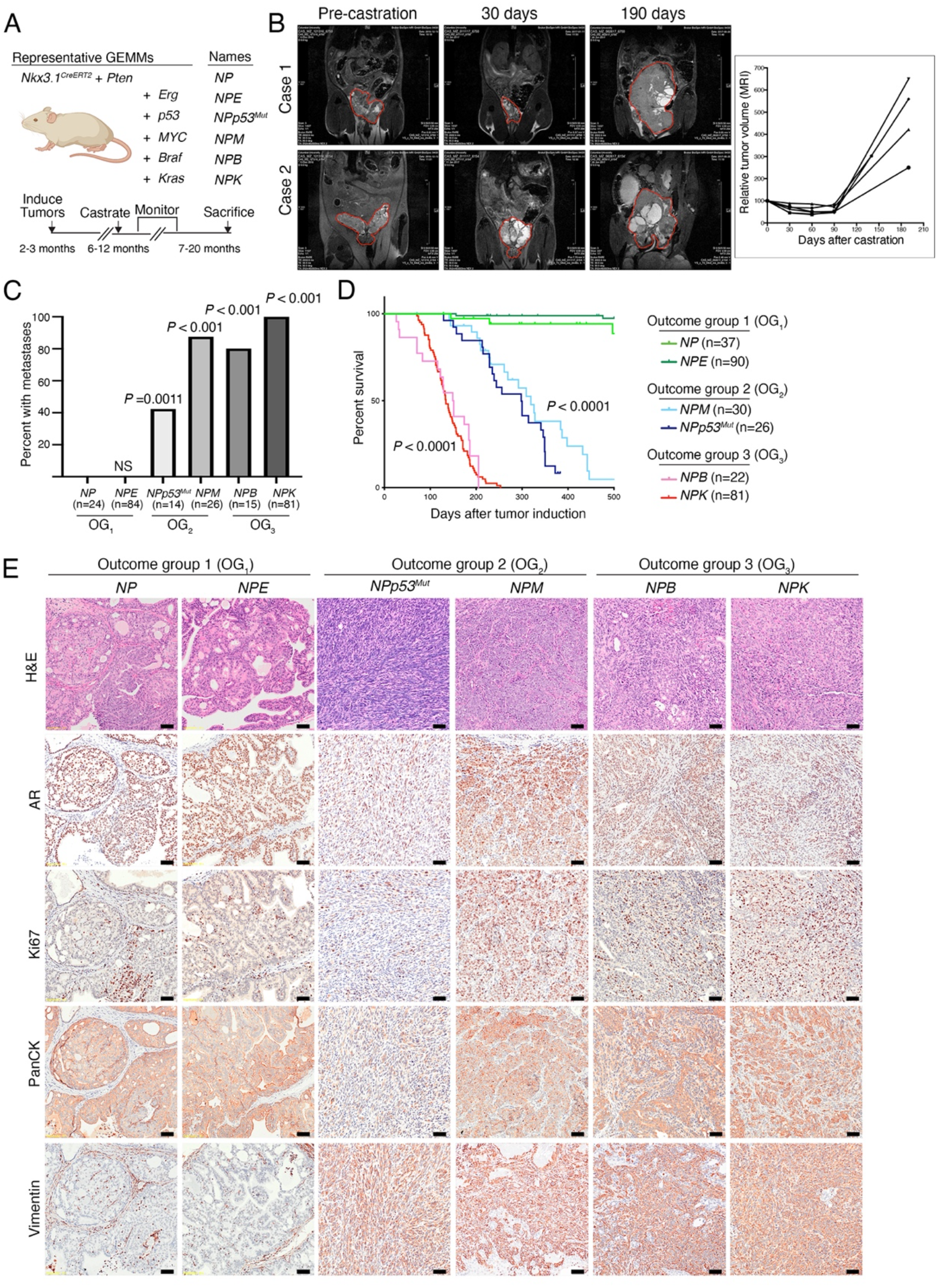
A GEMM resource that models prostate cancer progression. **A.** Schematic showing representative GEMMs used in this study. The GEMMs were generated by crossing *NP* mice (for *<u>N</u>kx3.1^CreERT2/+^; <u>P</u>ten^flox/flox^*) with the alleles shown in the panel to generate six complex strains (Table S1). The timeline for tumor induction, castration, monitoring, and sacrificing is shown at the bottom of the panel. **B.** MRI images showing tumor volume changes after castration of two representative *NPp53* mice (Case 1 and Case 2). The plot on the right shows the tumor volume changes over time for 4 representative *NPp53* mice. **C.** Frequency of metastasis observed in the GEMMs. The numbers of mice used to determine metastasis frequency for each model are indicated in parentheses; two-tailed *P*-values are shown for Fisher’s exact test comparing each model to the *NP* mice (control). OG, outcome group. **D.** Kaplan-Meier survival analysis showing three outcome groups (OG_1_, OG_2_, and OG_3_). *P*-values were calculated using a two-tailed log-rank test compared to the *NP* mice (control). For the analyses shown in (C) and (D), both intact and castrated mice were pooled for all GEMMs except *NPM*, where the effects of castration may be confounded by the AR-dependency of the Probasin promoter used to drive Myc expression (see Methods). **E.** Representative images for hematoxylin and eosin (H&E) (top row) and immunohistochemical staining of the indicated markers in primary tumors of the different GEMMs. Shown are representative images based on analyses 3 or more mice/group; scale bars represent 50 μm. See also Table S1, and Figures S1-S3.

These GEMMs are based on the *Nkx3.1^CreERT2^* allele (Fig. 2A) (35), which simultaneously introduces a prostate-specific inducible Cre driver and a heterozygous null allele for the *Nkx3.1* homeobox gene, as is common in early stage human PCa (Fig. S1A) (26). We generated the baseline *NP* mice (for *<u>N</u>kx3.1^CreERT2/+^; <u>P</u>ten^flox/flox^* mice) by crossing the *Nkx3.1^CreERT2^* allele with a *Pten* conditional allele (Fig. 2A) (36), since *PTEN* mutations are prevalent from the earliest to most advanced stages of human PCa (Fig. S1A) (25, 26). The *NP* mice were then crossed with various other alleles to model: *(i)* up-regulation of *Erg* in the *NPE* mice (*E* for *Rosa26^ERG/ERG^*, (37)); *(ii)* a missense mutation of *Tp53* in the *NPp53^mut^* mice (for *Trp53^LSL-R270H/flox^*, (38))*; (iii)* up-regulation of *c-Myc* in the *NPM* mice (*M* for *Hi-Myc*, (39)); *(iv)* an activating mutation of *Braf* in the *NPB* mice (B for *B-Raf^V600E^*, (40); and *(v)* an activating mutation of *Kras* in the *NPK* mice (*K* for *Kras^LSL-G12D^*, (41)) (Fig. 2A). These GEMMs also incorporate a conditionally-activatable fluorescent reporter, the *Rosa26-CAG-^LSL-EYFP^* allele (42), for high-efficiency lineage marking of tumors and metastases (30, 34).

Since tumor induction is based on an inducible Cre, expression of the relevant alleles following Cre-mediated gene recombination is not dependent on androgens, with the exception of the *Hi-Myc* allele, which is under the control of a constitutive *Probasin* promoter (39). Consequently, we could analyze tumor progression in both castration-sensitive and castration-resistant contexts (Fig. 2B-E, Fig. S2A). In particular, as evident by MRI imaging of *NPp53* mice, surgical castration leads to initial tumor regression followed by eventual outgrowth of castration-resistant tumors (Fig. 2B); therefore, this GEMM series recapitulates both hormone-sensitive and castration-resistant PCa.

This hormone-sensitive and castration-resistant GEMM cohort could be subdivided into three outcome groups (*OG*_1_ – *OG*_3_), based on their metastatic phenotype and overall survival (Fig. 2C-E; Fig. S2A,B; Table S1A,B). Those in *OG*_1_, which include the *NP* and *NPE* mice, developed indolent, non-lethal and non-metastatic tumors with mostly benign, PIN-like histology and low levels of proliferation, as evident by Ki67 staining. Those in *OG*_2_, including the *NPp53^mut^* and *NPM* mice, were characterized by lethality within one year, were prone to develop metastasis, and displayed highly heterogeneous and proliferative histopathological phenotypes. Finally, those in *OG*_3_, including the *NPB* and *NPK* mice, were characterized by lethality by 6 months of age, which was accompanied by highly penetrant metastasis and high-grade poorly differentiated histopathology.

To facilitate co-clinical investigations, we established allograft and organoid models from representative GEMMs (Fig. S3A,B, Table S1A). Allografts were generated by implanting freshly collected primary tumors into the flank of *nude* mouse hosts and passaged at least twice prior to analysis. Organoids were generated by FACS sorting to isolate the lineage-marked primary tumor cells, which were then cultured *in vitro* for up to five passages. The histopathology of the resulting allograft and organoid models were similar to the parental tumors from which they were derived (Fig. S3A,B). Interestingly, while we were able to generate organoids from GEMMs in each outcome group (*OG*_1_ – *OG*_3_), we were only able to generate allograft models from the *OG*_2_ and *OG*_3_ tumors, but not from indolent *OG*_1_ tumors. Thus, we have generated an extensive resource of PCa GEMMs, as well as established culturable and transplantable models of these GEMMs. Since the current study relies on co-clinical analyses *in vivo*, we therefore used the allograft rather organoid models; however, we envision that the organoid models will be beneficial for future *in vitro* investigations.

### GEMM subtypes recapitulate human PCa phenotypes

To characterize the molecular features of the GEMM cohort, we generated RNA-seq profiles from benign prostate tissue and prostate tumors from 136 individual mice. First, we reverse engineered a GEMM-specific ARACNe interactome from these RNA-seq profiles (Table S2A) (29). The resulting interactome outperformed our previously published mouse interactome, which was assembled from Illumina gene expression microarrays of a less comprehensive GEMM cohort (20). For example, bioactivity analysis—which assesses the ability of an interactome to recapitulate differential protein activity across distinct phenotypes (see Methods)—showed that differentially active proteins had significantly higher average normalized enrichment score (μ_NES,New_ = 3.79 in the new interactome, vs. μ_NES,Old_ = 1.96 in the previous one; *P* < 2.2×10^-16^, by 2-sided Kolmogorov– Smirnov test; Fig. S4A). For instance, while the androgen receptor (AR) regulon—a critical determinant of prostate differentiation and tumorigenesis (43)—was a poor predictor of AR activity in the previous interactome (20), it is highly predictive in the new one (see Fig. 3A, S4B,C).

**Figure 3:**
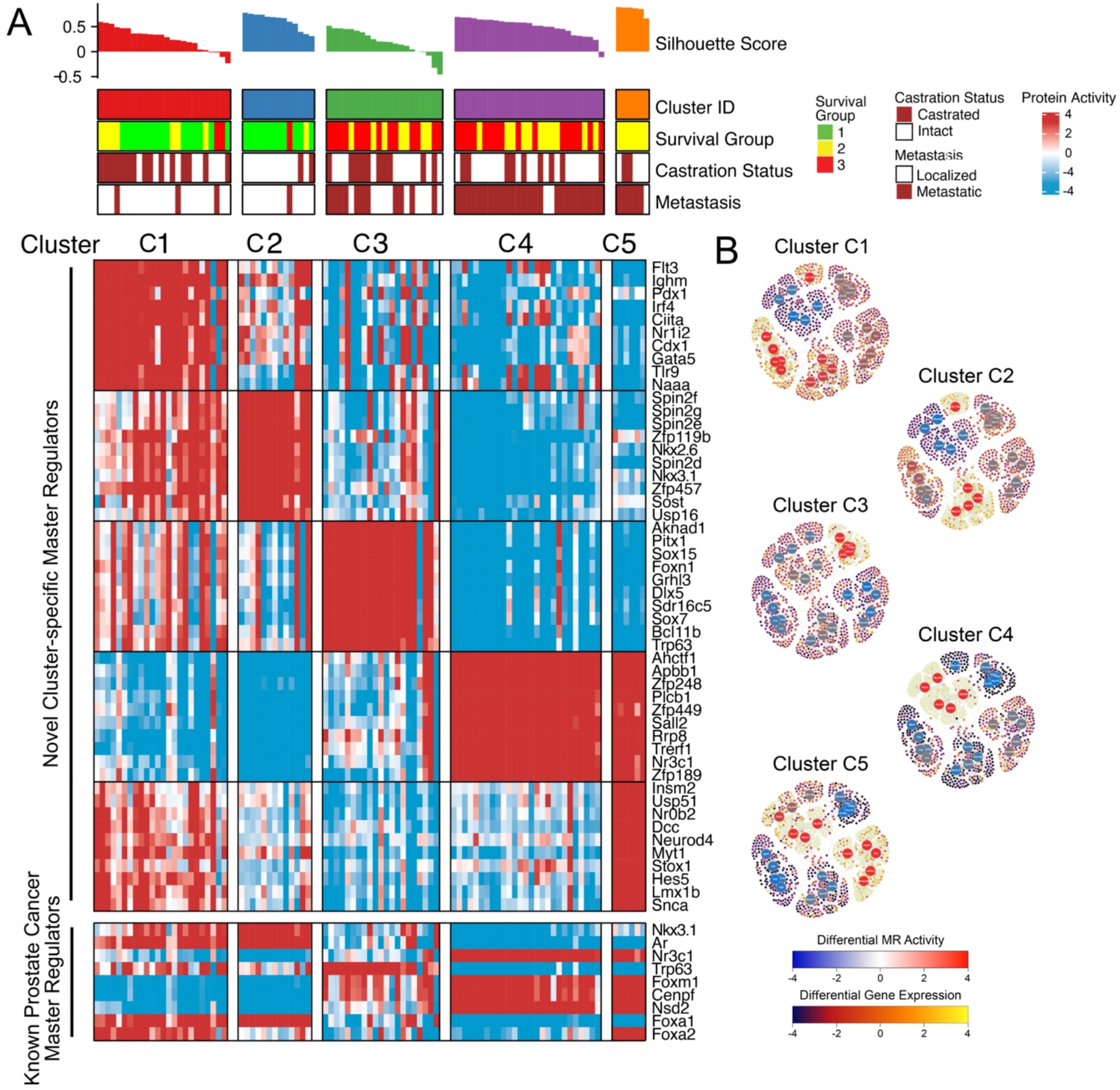
GEMM subtypes recapitulate human PCa phenotypes. **A.** Heatmap illustrating the results of protein activity-based cluster analysis of n = 91 GEMM-derived tumors (GEMM-DTs), as well as the silhouette score and correlative variables, such as survival group, castration status, and metastatic progression. This analysis identified five molecularly-distinct clusters (*C_1_ – C_5_*) that co-segregate with survival and metastatic potential. Shown for each cluster is the 10 most significantly activated MRs (top heatmap). Also shown are selected known human PCa markers (bottom heatmap). **B.** Representative sub-networks, representing the activity of the 25 most differentially active MR proteins (5/cluster, large circles) across all clusters and expression of their regulatory targets (small circles) on a cluster-by-cluster basis. Protein activity is shown using a blue (inactivated) to red (activated) scale, while target expression is shown on a blue (under-expressed) to yellow (over-expressed) scale. High resolution images with full visibility of the MRs is shown in Figure S5. See also Table S2, and Figures S4, S5.

To define GEMM molecular subtypes and to assess their relationship to human PCa, we focused on a subset of 91 GEMM-DTs having the most physiologically-relevant histopathological phenotypes (Table S1B). We first transformed their RNA-seq profiles to VIPER-measured protein activity profiles using the GEMM PCa interactome, and then performed protein activity-based cluster analysis (Fig. 3A; Table S2B, see Methods). To generate the differential gene expression signatures necessary for VIPER analysis, we compared each sample to the average of all 91 samples, thus identifying proteins with the greatest differential activity across all GEMM-DTs. Protein activity-based cluster analysis, which significantly outperforms gene expression-based clustering (22) identified 5 clusters, corresponding to 5 molecularly distinct subtypes (*C_1_* – *C_5_*) associated with disease aggressiveness with respect to outcome and metastasis (Fig. 3A; Fig. S4B; Table S2C), see Methods. In particular, subtypes *C*_1_ and *C*_2_ mainly comprise tumors from the indolent outcome group (*OG*_1_), while subtypes *C_3_* – *C_5_* comprise tumors from the two more aggressive outcome groups (*OG*_2_ and *OG*_3_). Interestingly, the more indolent subtype *C_1_* is mostly comprised of castration-resistant tumors while *C_2_* includes mostly hormone-sensitive tumors. Furthermore, > 90% of the tumors in *C*_4_ and *C*_5_ had progressed to metastases, compared to only 50% of those in *C*_3_, and 11% of those in *C_1_* and *C*_2_ (Fig. 3A; Fig. S4B).

With respect to their molecular phenotypes, *C*_1_ – *C*_5_ tumors were characterized by aberrant activity of a novel MR protein set that effectively distinguished each subtype (Fig. 3A). These include homeodomain proteins (e.g., Cdx1, Pdx1, Nkx2.6, Nkx3.1, Pitx1) and other transcriptional regulatory proteins (e.g., Gata5, Sox15, Sox7) that have known roles in cellular differentiation and cell lineage control in other tissue contexts. The coordinated switch between the molecular programs regulated by the top MR protein activities of each subtype was evident by the striking differential expression of the transcriptional targets of the most differentially active MRs of each subtype (Fig. 3B, Fig. S5).

While the top-most significantly active MRs in subtypes *C*_1_ – *C*_5_ are novel, differential activity of MR proteins with an established role in human and mouse PCa progression was clearly evident (Fig. 3A; Fig. S4C). In particular, the indolent subtypes *C*_1_ and *C*_2_ present high activity of Nkx3.1, p63, and Ar, which are associated with well-differentiated PCa (43). In contrast, the most aggressive and highly metastatic subtypes *C_4_* and *C_5_*, present high activity of FoxM1, Cenpf, and Nsd2, which are all aberrantly activated and functionally necessary for aggressive PCa, in both humans and mice (20, 44), as well as activation of the glucocorticoid receptor (Nr3c1), which is activated in human CRPC (45). Interestingly, subtype *C*_5_ presented dysregulation of proteins associated with NEPC, including Foxa2 and Foxa1 (Fig. 3A, Fig. S4C) (46) and was significantly enriched in the Beltran NEPC signature (*P* < 10^-4^, see Methods) (11), while no enrichment for the NEPC signature was detectable in subtypes *C*_1_ – *C*_4_. Notably, none of the GEMM-DTs, including those associated with advanced PCa and NEPC (*C_4_* and *C_5_*, respectively), had undergone pharmacological treatment; this is a novel feature of the GEMM cohort compared to analogous human PCa cohorts, which are generally derived from patients that had undergone extensive treatments (e.g., (25)).

To further characterize the GEMM subtypes and to assess their relationship to human PCa, we performed pathway enrichment analyses using > 900 relevant gene sets, collected by the Broad Institute MSigDB (47), Kyoto Encyclopedia of Genes and Genomes (KEGG) (48) and REACTOME (49) collections (Table S2D). To determine the top pathways associated with each subtype, we computed the enrichment of each gene set in proteins differentially active in each sample, and then integrated across samples in each cluster using Stouffer’s method (see Methods). These analyses revealed that the most aggressive subtypes, *C*_4_ and *C*_5_, show strong enrichment for proliferation and oncogenic hallmarks, e.g., G2M Checkpoint (*P* = 2.14×10^-39^), E2F Targets (*P* = 4.5×10^-88^), DNA Repair (*P* = 5.8×10^-68^), and MYC Targets V1 and V2 (integrated *P* = 3.3×10^-21^) (Fig. S4D; Table S2D). (Note that all *p*-values reported in this manuscript are corrected for multiple hypothesis testing, see Methods). Interestingly, interferon and inflammatory response hallmarks, including interferon alpha response (*P* = 2.4×10^-52^), interferon gamma response (*P* = 8.2×10^-68^) and inflammatory response (*p* = 4.7×10^-18^) were downregulated in subtypes *C*_4_ and *C*_5_ (Fig. S4D; Table S2D), suggesting that the most aggressive tumors may have an immunosuppressive tumor microenvironment.

Taken together, these molecular analyses define a series of GEMM subtypes (*C_1_* – *C_5_*) that: (i) are distinguished by novel as well as known MR protein activities; (ii) model a wide range of PCa phenotypes from indolent to aggressive variants, including castration-sensitive and castration-resistant tumors; and (iii) can differentiate lethal subtypes of adenocarcinoma and NEPC in the absence of prior treatment. Thus, this GEMM cohort represents a valuable resource to characterize and model human PCa, particularly for advanced tumors, as would be the principal focus for predicting drug treatments for human patients.

### Matching GEMM-DTs to patient tumors and metastases

Having established that the molecular programs of the GEMM cohort are relevant for human PCa, we next asked whether individual GEMM-DTs can provide high-fidelity (cognate) surrogates for individual patients (Fig. 4). We thus compared the MR protein activity signature for individual human patients with that of each available GEMM-DT (n = 91) and designated those presenting highly significant conservation of patient specific MR activity (*P* ≤ 10^-5^, by 1-tail aREA test) as cognate models. For these studies, we queried two well-characterized patient cohorts, one comprised of treatment naïve primary tumors collected by The Cancer Genome Atlas (TCGA, n = 333) (26), and a second comprised of post-treatment metastatic biopsies from mCRPC patients collected as part of the Stand Up to Cancer-Prostate Cancer Foundation cohort (SU2C, n = 212) (25).

**Figure 4:**
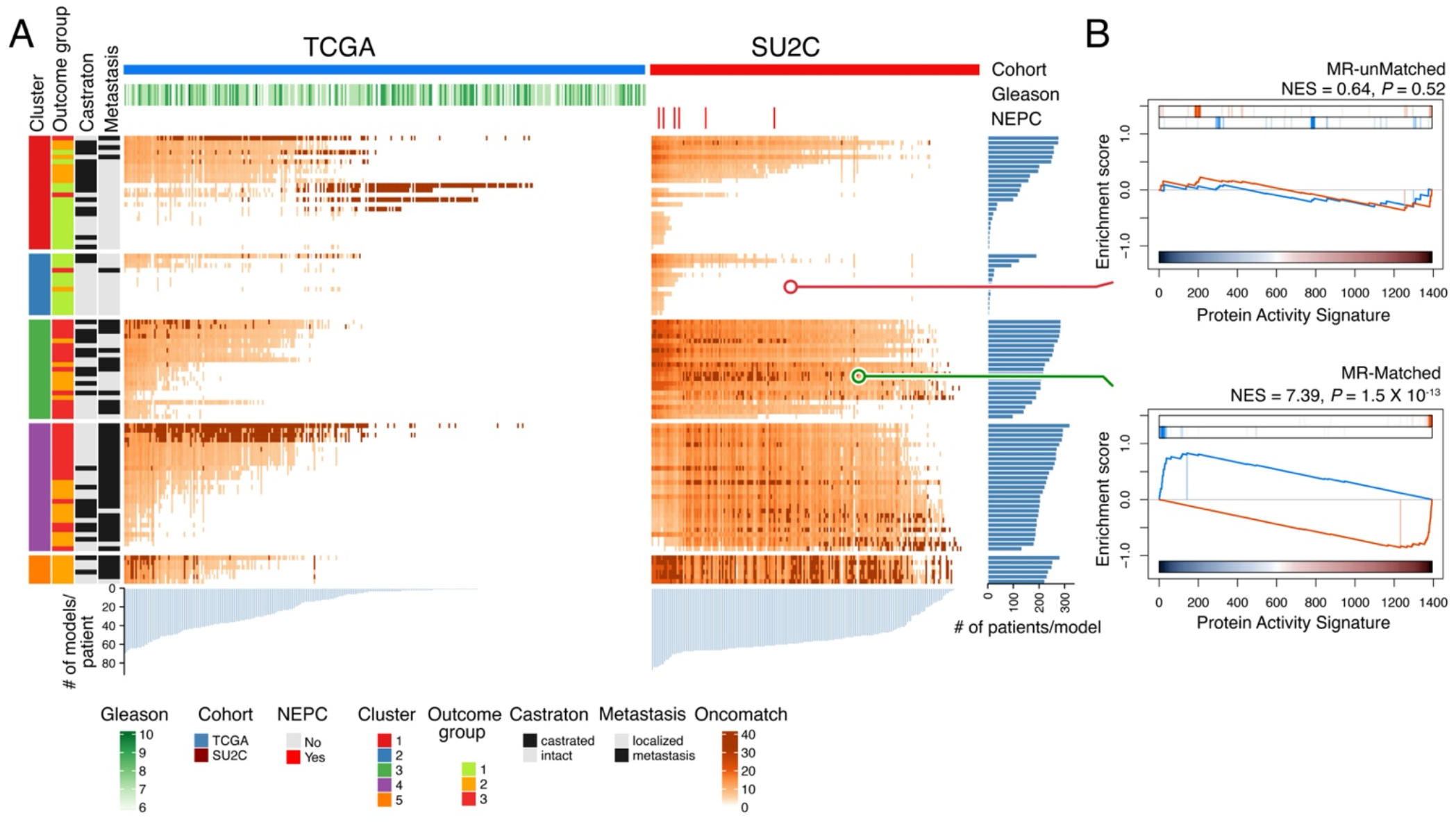
Matching GEMM-DTs to patient tumor and metastases. **A.** Heatmaps representing the MR-based fidelity score of each patient (columns) versus each GEMM-DT model (rows), as assessed for TCGA (right) and SU2C (left) patients, respectively. Relevant patient phenotypic variables—*i.e.*, Cohort, Gleason score, and NEPC status—are shown in the top three bars, while relevant GEMM-DT phenotypic variables—*i.e.*, Cluster, outcome, castration status, and metastasis status—are shown in the four vertical bars to the left of the heatmap. Fidelity scores are computed as the –Log_10_*P* of the patient vs. GEMM-DT MR enrichment analysis. The 5 top-most significant cognate models for each patient are shown in dark red; other statistically significant (P ≤ 10^-5^) high-fidelity models are shown using a lighter to darker color scale (as shown). The light blue barplots at the bottom of the two heatmaps shows the number of candidate cognate models for each patient, while the dark blue barplots to the right show the number of patients for which a GEMM-DT represents a cognate model. Overall, 78% of the samples in the TCGA (n = 261 of 334) and 93% of those in the SU2C cohorts (n = 198 of 212) have at least one cognate GEMM-DT. **B.** GSEA of the fidelity analysis for representative GEMM-DT-SU2C pairs showing an example of an MR-unmatched (low-fidelity, top) and an MR-matched (high-fidelity, bottom) pair. See also Tables S3, S4.

First, we assembled distinct human PCa interactomes for the primary tumors and metastases by ARACNe analysis of RNA-seq profiles from the TCGA and SU2C cohorts, respectively (Table S3A,B, see Methods). We then used VIPER to transform the transcriptional profiles of the TCGA and SU2C cohorts into protein activity profiles, using their respective interactomes (Table S3C,D). Differential expression signatures for VIPER analysis were computed using as reference a combination of all RNA seq profiles from the GTEx normal prostate cohort (n = 245) (50) and a humanized version of normal mouse prostate tissue (see Methods). This approach allowed identification of MR proteins that are specifically dysregulated in the tumor context compared to normal prostate.

To identify cognate models on an individual patient basis, we generated protein activity signatures by comparing each GEMM-DT to the same reference as above. We then assessed the fidelity of each of the GEMM-DTs to each individual TCGA or SU2C patient by assessing the enrichment of the 25 most activated (25↑) and 25 most inactivated proteins (25↓) of each human tumor, representing its *candidate MR proteins* (MR proteins hereafter for simplicity), compared with MR proteins differentially active/inactive in each GEMM-DT (Fig. 4; Table S4A,B). We used fixed number of MR proteins (i.e., 25↑+25↓) since: (i) this is required to make the statistics of enrichment analyses comparable across proteins and cohorts; and (ii) we have previously shown that an average of 50 MRs is sufficient to account for the canalization of functionally-relevant genetic alterations in the vast majority of TCGA samples (22). For simplicity we refer to these 25↑+25↓ MR proteins as the MR-activity signature. Having assessed enrichment of their MR-activity signatures, we selected a highly conservative statistical threshold (*P* ≤ 10^-5^) to nominate high-fidelity, cognate GEMMs-DTs, see Methods.

Analysis of primary tumors from the TCGA cohort revealed that 78% had at least one high-fidelity cognate GEMM-DT (n = 261/334; Fig. 4; Table S4A). Strikingly, analysis of PCa metastases from the SU2C cohort revealed an even greater fraction (93%) of patients with high-fidelity cognate GEMM-DTs (n = 198/212) and, on average, 48 cognate GEMM-DTs were identified as significant cognate models (*P* ≤ 10^-5^) for each SU2C tumor (Fig. 4; Table S4B). This likely reflects the inherent bias of the GEMM cohort towards more aggressive PCa phenotypes (see Fig. 3). Heatmap representation of the matched patients and GEMM-DTs shows good clustering of patient and GEMM-DTs in the same subtype (*C*_1_ – *C*_5_). While most patients were matched to multiple GEMM-DTs, we highlight the GEMM-DTs representing the top 5 most statistically significant MR-based matches for each patient (i.e., the highest-fidelity models, Fig. 4), since these would provide the most realistic models for co-clinical studies. As expected, given the more aggressive nature of the SU2C patients, highest-fidelity models for the TCGA and SU2C cohorts formed distinct clusters. These findings demonstrate that individual tumors from the GEMM cohort (GEMM-DTs) represent high-fidelity surrogates for individual human PCa patients. Furthermore, most of the patients with aggressive PCa, who would benefit most from co-clinical validation of novel treatments *in vivo*, are represented by at least one GEMM-DT.

### Generation of drug perturbational profiles to identify MR-inverter drugs

As discussed above, two PCa cell lines—the androgen-dependent LNCaP and the androgen-independent DU145 cell lines—jointly provided high-fidelity models for >80% of the tumors in the TCGA cohort, based on MR-activity signature conservation (27). To predict optimal treatments for PCa patients with aggressive tumors, we focused on the more aggressive DU145 cell line. Specifically, drug MoA was assessed from RNA-seq profiles of DU145 cells harvested at 24h following treatment with 117 FDA-approved and 218 late-stage experimental (i.e., in Phase II and III clinical trial) drugs, as well as vehicle control (DMSO) (i.e., n = 335 drugs in total, Table S5A). To minimize activation of cell death or cellular stress pathways that would confound assessment of drug MoA, cells were treated with the 48h EC_20_ concentration of each drug (i.e., highest sublethal concentration), as assessed from 10-point drug response curves (see Methods). RNA-seq profiles were generated using PLATE-Seq, which was specifically designed to generate profiles of drug perturbations (51).

For each drug, differential protein activity profiles, representing the drug’s MoA, were then generated by VIPER analysis of drug vs. vehicle control-treated cells (Table S5B, see Methods). Cluster analysis of differentially activated proteins following drug treatment with the most bioactive agents—*i.e.*, 115 drugs inducing the most significant differential protein activity patterns— identified 11 drug clusters (D_1_ – D_12_) based on differential activation/inactivation of 13 protein sets (*programs*) (P_1_ – P_13_) (Fig. S6). Consistent with the analysis, drugs presenting similar MoA were clustered together or in closely related clusters. For instance, cytotoxic drugs cosegregated in clusters D_1_, D_2_, D_5_, D_7_ and D_8_, while kinase inhibitors were found mostly in clusters D_3_, D_10_, and D_12_, including a majority of MAPK (*i.e.*, sorafenib, dabrafenib, vemurafenib, trametinib), EGFR inhibitors (*i.e.*, lapatinib, erlotinib, vandetanib), and mTor inhibitors (*i.e.*, everolimus, temsirolimus) in D_12_. Similarly, a subset of hormone blockade drugs (i.e., anastrazole, enzalutamide, abiraterone, mitotane) clustered in D_6_ and D_9_, and another subset (i.e., raloxifene, leuprolide, exemestane, tamoxifen) in D_3_ and D_4_. Folate (*i.e.*, methotrexate, pralatrexate) and microtubule inhibitors (i.e., paclitaxel, ixabepilone) clustered in D_4_, while proteasome inhibitors (i.e., ixazomib, carfilzomib, bortezomib) presented very similar profiles in D_11_ and D_12_. In contrast, as expected, more pleiotropic drugs with broad-spectrum MoA, such as HDAC inhibitors (*i.e.*, vorinostat, panobinostat, belinostat), CRBN inhibitors (*i.e.*, thalidomide, lenalidomide, pomalidomide), and de-methylating agents (*i.e.*, decitabine, azacytidine), were broadly distributed across multiple clusters. Given the high reproducibility of replicate perturbational profiles from the same drug (*P* ≪ 0.05, by 2-tail enrichment analysis), for the majority of the 115 bioactive compounds, and the diversity of the differential protein activity they induce, this suggests that the DU145 perturbational profiles effectively inform on drug MoA.

### Using OncoLoop to predict candidate drugs for individual patients

To predict candidate drugs, we identified those for which the MR-activity signature of both a patient and its cognate GEMM-DT was significantly inverted in drug-treated vs. vehicle control-treated cells (MR-inverter drugs) (Fig. 5A-C). Since our goal is to identify treatments for patients with advanced, rather than indolent, PCa, we focused on the metastatic patients in the SU2C cohort and their cognate GEMM-DTs (which could be identified for 93% of the SUC2 patients; see Fig. 4). For each SU2C patient and each cognate GEMM-DT, MR-inverter drugs were identified by enrichment analysis of the respective top 25↑ and 25↓ MR proteins (see Fig. 4) compared with the MR activity signatures of the drug-treated vs. vehicle control-treated cells, at a conservative statistical threshold (*P* ≤ 10^-5^) (Table S5C,D).

**Figure 5:**
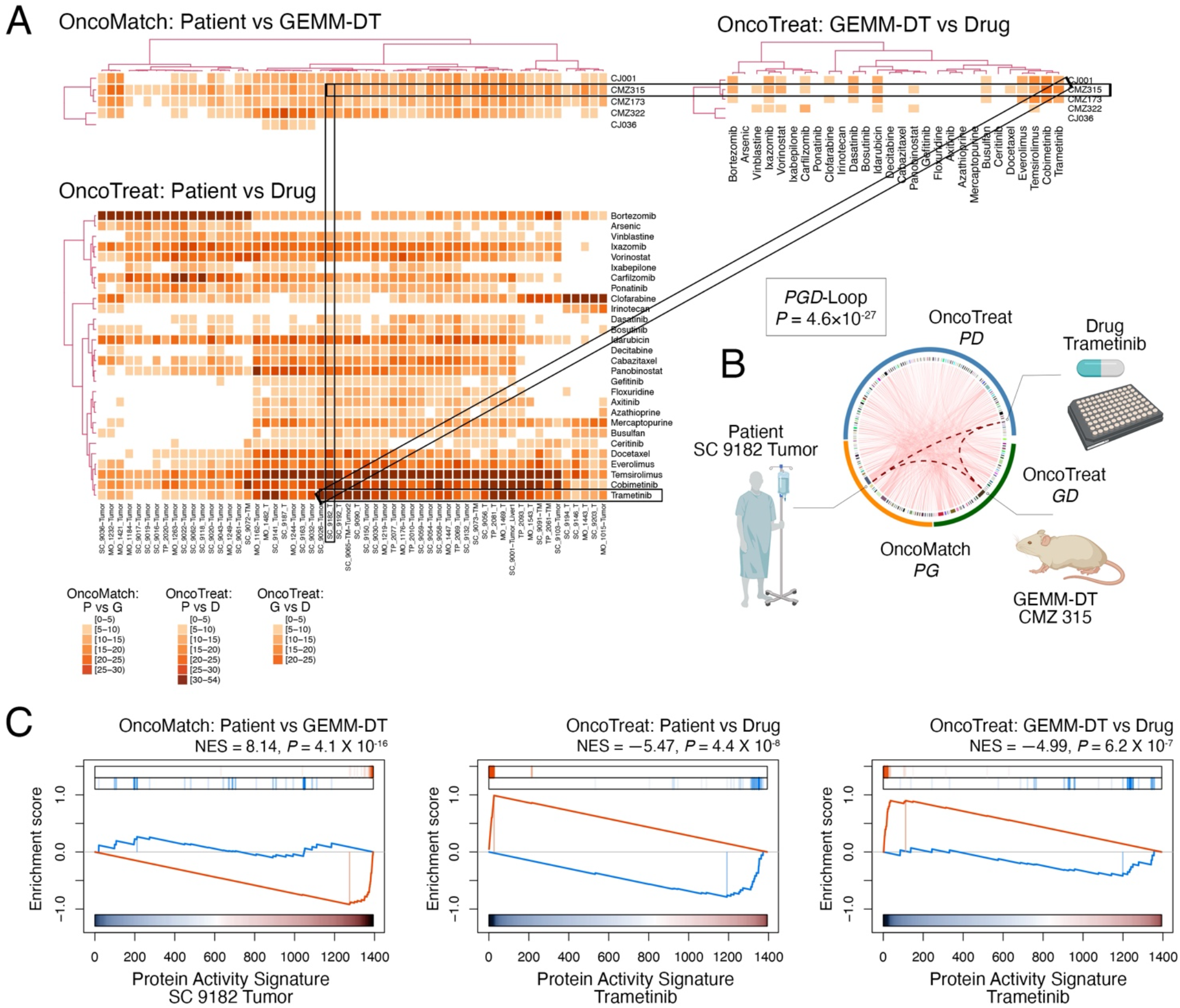
OncoLoop analysis. **A.** Examples of a PGD-Loops, showing three heatmaps for a representative subset of patients, GEMM-DTs, and drugs: the top left heatmap (OncoMatch: Patient vs. GEMM-DT) depicts the fidelity scores for 56 SU2C samples (columns) and 5 GEMM-DTs (rows); the bottom left heatmap (OncoTreat: Patient vs. Drug) shows the MR-inverter scores for 28 drugs (rows) as assessed against 56 SU2C samples (columns); and the top right heatmap (OncoTreat: GEMM-DT vs. Drug) shows the MR-inverter scores for the same 28 drugs (columns) as assessed against the 5 GEMM-DTs (rows). All scores are computed as (−Log_10_*P*) and statistically significant scores (P ≤ 10^-5^) are shown with a light to dark color scale, as indicated; non-significant scores are shown in white. MR-inverter scores are based on MR activity inversion analysis based on the drug-vs. vehicle control-treated DU145 cells. For visualization purposes, heatmap results are depicted by hierarchical clustering. One among many statistically significant PGD-Loops is highlighted by the rectangles. **B.** Circos plot showing the PGD-loop highlighted in panel A, comprising a SU2C patient (SC_9182_T), his top-ranked cognate GEMM-DT (CMZ315), and the drug trametinib; the *P-*value was calculated by integration of there associated scores. **C.** The corresponding GSEA plots for the PGD-loop in Panel B, showing: the patient to cognate GEMM-DT fidelity analysis (OncoMatch: Patient vs. GEMM-DT, left); the MR-inversion score by trametinib, as assessed for the patient’s MRs (OncoTreat: Patient vs. Drug, middle), and the MR-inversion score by trametinib, as assessed for the cognate GEMM-DT (OncoTreat: GEMM-DT vs. Drug, right). See also Tables S5, S6, and Figure S6.

The resulting PGD-loops—comprising a patient, its cognate GEMM-DT, and the candidate MR-inverter drugs (Fig. 5A)—were ranked based on the Stouffer’s integration of three *z*-scores, corresponding to: (i) *z_PG_*, the GEMM vs. patient MR-activity conservation *z*-score (Table S4B); (ii) *z_PD_*, the patient-specific MR-inverter drug *z*-score (Table S5C); and (iii) *z_GD_*, the cognate GEMM-DT-specific MR-inverter drug *z*-score (Table S5D). Considering all possible combinations of SU2C patients (n = 212), GEMM-DTs (n = 91), and drugs (n = 337), there were >6.5M potential PGD-loops; of these 668,138 achieved statistically significance (*P* ≤ 10^-5^) on all three *z*-scores (Table S6). Notably, the extensive coverage of both GEMM-DTs and drugs for the SU2C patients, which on average was 48 cognate GEMM-DTs (*P* ≤ 10^-5^) for each SU2C tumor (see Fig. 4) and 22 FDA-approved candidate MR-inverters (*P* ≤ 10^-5^) for each SU2C tumor/cognate-GEMM-DT pair, supports the use of OncoLoop in a co-clinical setting.

To provide a visual representation of OncoLoop, we show three heatmaps reporting the statistical significance of model fidelity (OncoMatch), and drug MR-inversion analyses (OncoTreat) for a subset of SU2C patients (n = 56), their cognate GEMM-DTs (n = 5), and drugs (n = 28), which were especially enriched in statistically-significant PGD-Loops (Fig. 5A). For illustrative purposes, we highlight a PGD-loop defined by an mCRPC patient (SC_9182_T), its cognate GEMM-DT (CMZ315)—harboring mutation of p53, a gene that is frequently dysregulated in human CRPC (see Fig. S1A)—and the drug trametinib, which is a MEK inhibitor (Fig. 5A,B). Notably, as evident by GSEA, all three Normalized Enrichment Score (NES) values for this PGD-Loop were highly significant, including for the model fidelity analysis (NES = 8.14, *P* = 4.1×10^-16^), as well as for the patient-specific (NES = −5.47, *P* = 4.4×10^-8^), and cognate GEMM-DT-specific (NES = −4.99, *P* = 6.2×10^-7^) trametinib-mediated MR-inversion (Fig. 5C), resulting in a highly significant integrated *p*-value (NES = 10.71, *P* = 4.6×10^-27^).

### Co-clinical validation in cognate GEMM-DTs

To optimize the clinical translation of this approach and to capture Oncoloop predictions for a majority of patients, we focused on PGD-Loops comprising drugs that were both clinically-available and most recurrently predicted for SU2C patients (Fig. 6A). Specifically, we considered only FDA approval drugs (n = 117), predicted for ≥ 50% of the SU2C tumors, and active at physiologically-relevant concentrations (≤ 1μM), which prioritized 16 candidate drugs for validation. Cluster analysis of their MR-inversion *z*-score for individual SU2C tumors (i.e., predicted drug sensitivity) identified 5 clusters, which represent subsets of patients predicted to be sensitive to the same drugs (*i.e.*, *pharmacotypes*; see Methods) (Fig. 6A). Concurrent, yet fully independent, stratification of the GEMM-DTs, also considering only FDA approval drugs active at physiologically-relevant concentrations (≤ 1μM), identified the same top 16 drugs (Fig. 6B), thus further confirming the representative nature of these cognate models.

**Figure 6:**
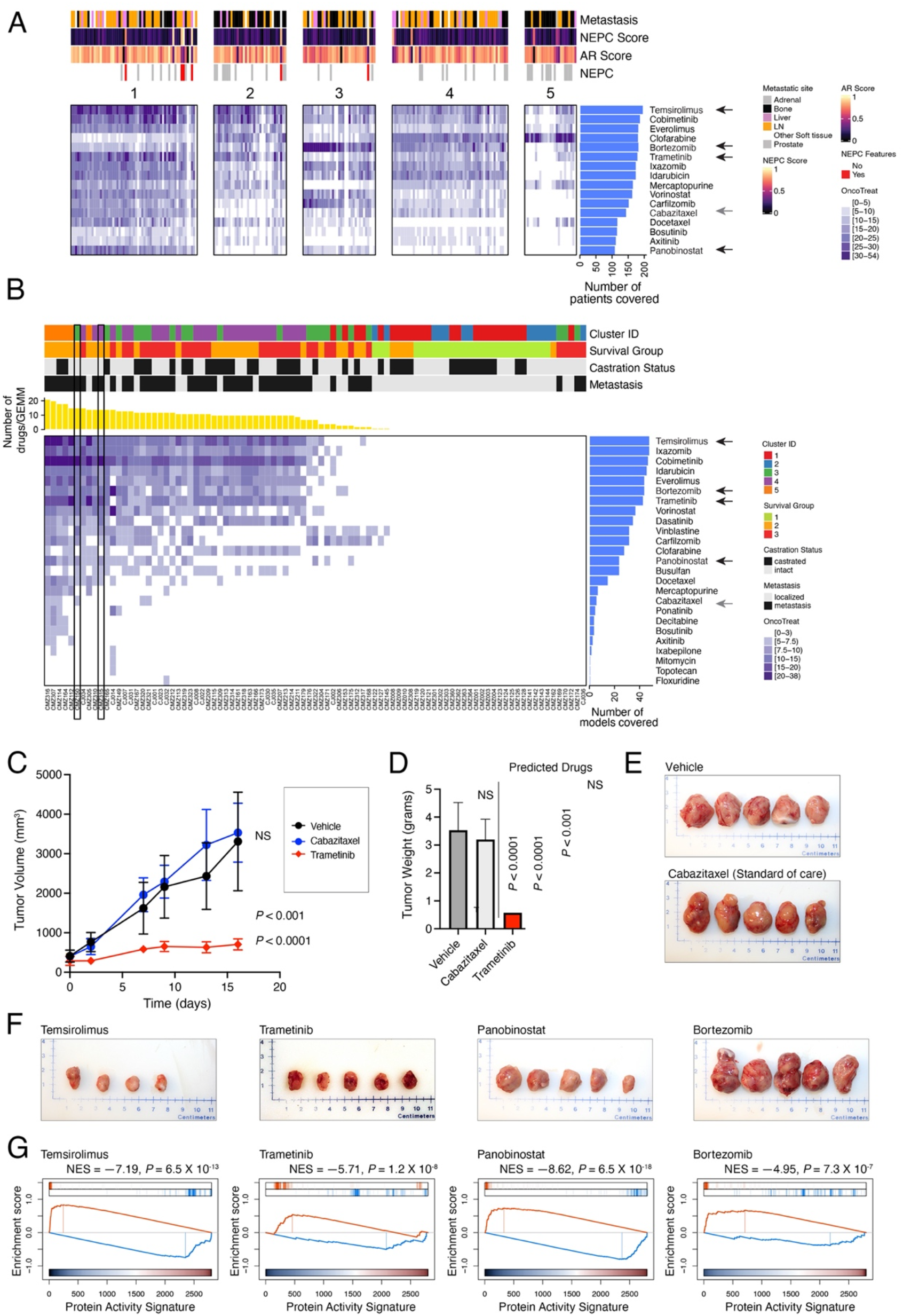
Selection and validation of candidate drugs. **A,B.** <u>Drug prioritization</u>: FDA approved drugs (n = 117) (rows) were prioritized as candidate MR-inverters of either patients from the SU2C cohort (n = 212) (columns in Panel A) or GEMM-DTs (n = 91) (columns in Panel B), using drug perturbation data from the DU145 cells. Drugs were filtered based on screened concentration (≤ 1uM) and patient coverage, *i.e.*, only those identified as a MR-inverters for >50% of the human samples were considered. Relevant phenotypic variables for either patients or GEMM-DTs are shown in bars at the top each heatmap. The MR-inverter score (−Log_10_*P*, as computed by aREA) is shown using a white (*P* > 10^-5^) to dark blue heatmap (see legend). The blue barplots on the right summarize the number of patients or GEMM-DT predicted as sensitive to each drug. Black arrows to their right point to candidate drugs selected for validation, while the grey arrows point to cabazitaxel, the standard-of-care for mCRPC, which was significant in the patient but not in the GEMM-DT analysis. In panel B, the yellow barplot at the top shows the number of drugs identified as significant MR-inverters for each GEMM-DT and the rectangle indicates the allografts used for validation. See also Figure S7. **C-F.** <u>Validation in the allograft model</u>. Selected drugs were validated *in vivo*, in allograft models derived from the cognate GEMM-DT CMZ315. Allografts were grown subcutaneously in *nude* mouse hosts and the mice were treated with predicted drugs, vehicle control, and a negative control (cabazitaxel) for the times indicated. **C.** Summary of tumor volume changes over the treatment period. **D.** Summary of tumor weights, following sacrifice. *P*-values for C and D were computed by one-way ANOVA at the last time point, compared to Vehicle treated tumors and adjusted for multiple hypothesis testing with Dunnett’s test (10 animals were enrolled to the vehicle control arm and 5 animals were enrolled on each of the drug treatment arms). **E.** Representative images of final tumor sizes in vehicle control and negative control-treated allografts. **F.** Representative images of final tumor sizes in allografts treated with predicted drugs. **G.** Pharmacodynamic assessment of MR-inversion by GSEA analysis for the four predicted drugs comparing drug-mediated of the drug-versus vehicle-treated tumors.

Among these 16 drugs, we eliminated those with overlapping MoA (e.g., multiple HDAC inhibitors) or lacking demonstrated variable efficacy in prior PCa clinical trials, which yielded four candidates for experimental validation: temsirolimus (m-TOR inhibitor) (NCT00919035, NCT00012142) (52); trametinib (MEK inhibitor) (NCT02881242, NCT01990196), panobinostat (HDAC inhibitor) (NCT00667862, NCT00878436) (53), and bortezomib (proteosome inhibitor) (NCT00193232, NCT00183937). As a negative control, we selected cabazitaxel—a current standard of care for advanced PCa—which was predicted as a significant MR-inverter for the human tumors but not for their cognate GEMM-DTs (see Fig. 6B).

To test drug sensitivity predictions, we performed tumor growth assays in allografts derived from two cognate GEMM-DTs, namely CMZ315 and CMZ150 (Fig. 6C-F; Fig. S7A-D). These represent two of the aggressive GEMM clusters, *C*_3_ and *C*_4_, and were derived from p53-mutated and MYC-amplified mCRPC GEMM tumors, respectively (Table S1). For each drug, we used their published conditions for *in vivo* mouse studies to determine their appropriate concentration and treatment schedule, and we also confirmed their uptake into allograft tumors, by pharmacokinetic assays (Fig. S7A, see Methods).

Three of the four predicted drugs—namely temsirolimus, trametinib, and panobinostat— significantly reduced tumor volume and weight in allografts from both of the cognate GEMM-DTs (*P* ≤ 0.01, one-way ANOVA, Fig. 6C-F; Fig. S7B-D). Bortezomib did not significantly inhibit tumor growth in either allograft, potentially due to cell adaptation mechanisms that buffered the initial MR-inversion. Also consistent with predictions, cabazitaxel induced only modest tumor volume/weight reduction, which was significant in only one of the two allografts. Taken together, these findings confirm that drugs predicted via OncoLoop to mediate MR-inversion are frequently capable of abrogating tumor growth when experimentally validated *in vivo*.

Lastly, we performed pharmacodynamic studies to assess the ability of the predicted drugs to recapitulate MR-inversion (Fig. 6G). To assess MR-activity inversion before significant tumor cell death or necrosis could ensue, we analyzed RNA-seq profiles from short-term (5-day) drug-vs. vehicle control-treated allograft (CMZ315). Confirming OncoLoop predictions, all four candidate drugs induced highly significant MR-inversion *in vivo*, as evidenced by GSEA analyses showing significantly negative NES values (*P* < 10^-7^, for all tested drugs, by 1-tail aREA test; Fig. 6G).

### Co-clinical validation in cognate human PDX models

To validate OncoLoop predictions in human tumor contexts, we performed analogous co-clinical studies using the well-characterized LuCaP series of PDX models, which were established from primary tumors and metastases obtained from University of Washington Tissue Acquisition Necropsy program (15). Notably, the LuCaP PDX models were developed from biologically heterogeneous advanced PCa tissues from primary and metastatic sites and include a range of tumors that vary in their response to castration, with some displaying castration-sensitivity (15).

To identify cognate PDX models, we first generated RNA-seq profiles for 5 LuCaP PDX models that had been perturbed with multiple drugs and used these RNA-seq profiles to generate a PDX interactome (Table S7A, see Methods). Among all 5 baseline LuCaP PDX models, VIPER analysis identified LuCaP-73 as the most significantly matched to the GEMM-DTs used herein (Fig. 7A). Furthermore, among the candidate drugs identified for the human tumors (see Fig. 6A), several were identified as significant MR-inverters for the LuCaP-73 PDX, including trametinib and panobinostat (Fig. 7B; Table 7B). We therefore tested whether these two drugs could abrogate LuCaP-73 viability *in vivo.* Indeed, both trametinib and panobinostat showed near-complete tumor growth inhibition (*P* < 0.0001; Fig. 7C-E). These findings provide important support for the translation of OncoLoop to a human PCa context.

**Figure 7:**
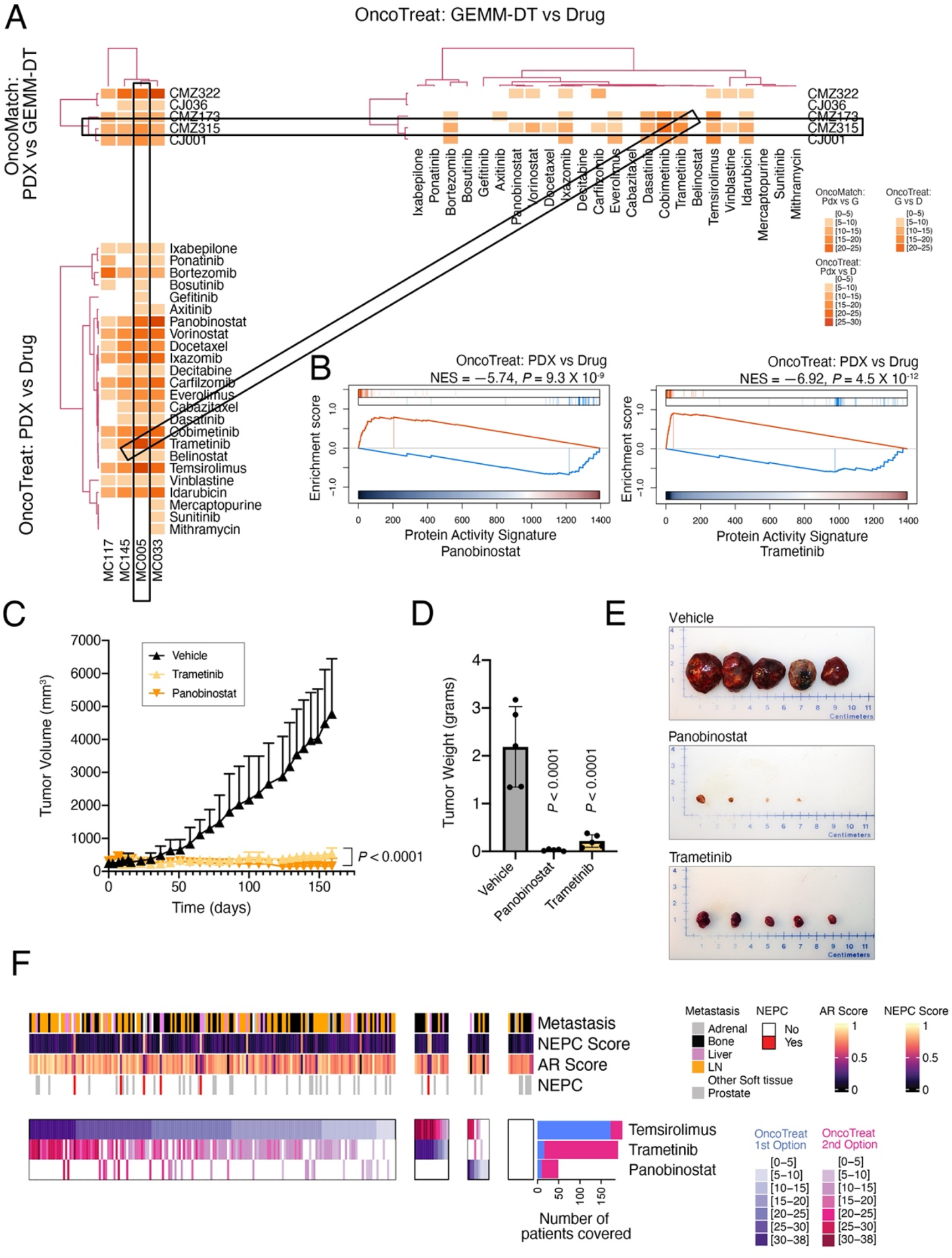
Co-clinical validation of candidate drugs. **A, B.** <u>OncoLoop analysis of patient derived xenograft (PDX) models</u>: **A.** Similar to Figure 5A, shown are three heatmaps representing the fidelity and MR-inverter scores for 4 LuCaP PDX tumors (columns in left heatmaps), 5 GEMM-DTs, and 28 drugs. The rectangles show a representative PGD-Loop, comprising a PDX (MC005/LuCaP73), its cognate GEMM-DT (CMZ315), and two of the top-predicted drug candidates evaluated in the allograft models (panobinostat and trametinib). For visualization purposes, heatmaps were clustered as in Figure 5A. **B.** GSEA showing the MR-inverter *P*-value for panobinostat and trametinib based on the MC005/LuCaP73 model. **C-E.** <u>Validation in the PDX model.</u> The MC005/LuCaP73 PDX was grown in *nude* mouse hosts and treated with predicted drugs or vehicle for the times indicated. **C.** Summary of changes in tumor volume over the treatment period. **D.** Summary of tumor weights following sacrifice. *P*-values for C and D were computed by one-way ANOVA at the last time point, compared to vehicle control-treated models and adjusted for multiple hypothesis testing with Dunnett’s test. **E.** Representative images of final tumor sizes. **F.** <u>Pharmacotype analysis</u>: Identification of patient subsets predicted to elicit sensitivity to the same drug subsets, by cluster analysis. Shown are four subtypes identified, including patients most sensitive to temsirolimus, trametinib, panobinostat, as well as patients for which none of the three validated drugs was significant. For each patient, the score of the most statistically significant MR-inverter drug is shown using a white (non-significant) to dark-blue color map, see legend; the second most significant drug is shown using a white (non-significant) to dark-red color map, see legend. This heatmap provides the rationale for a possible umbrella or combination trial where each patient (column) could be randomized to the most statistically significant MR-inverter drug (rows) validated in the preclinical study, based on its MR-inversion score, or to the combination of the two most significant drugs. The barplots to the right show the total number of patients predicted to be sensitive to each drug, as either most significant (blue) or second best selection (red). See also Table S7.

## Discussion

Predicting individualized drug efficacy in human patients remains a key challenge in precision medicine, and relatively few approaches have been described to identify models that best recapitulate patient-relevant drug response (6). To address these challenges, we have developed OncoLoop, which uses quantitative protein activity-based metrics to first identify high-fidelity and preexisting models for individual patient’s tumors, then to predict drug efficacy for a given patient’s tumor and its cognate model, and lastly to validate the drug predictions in the preexisting cognate model. In the current study, we demonstrate the effectiveness of OncoLoop in the context of PCa. In particular, by leveraging an extensive and diverse collection of GEMMs, we find that > 90% of mCRPC patients in a published PCa cohort are represented by at least one cognate GEMM-DT. We then use large-scale drug perturbation data from a context-matched PCa cell line to predict MR-inverter drugs for the patients and their cognate models. The predicted MR-inverter drugs were experimentally validated both in allografts from the GEMM-DT as well as in a cognate PDX model. Notably, this approach can be applied to evaluate candidate drugs for clinical trials, including umbrella and basket trials, based on *in vivo* validated drug stratification across models representing distinct pharmacotypes (e.g., Fig. 7F). Thus, OncoLoop represents an accurate and versatile framework for both predicting and evaluating individualized drug treatments in real-time. Notably, since all of the resources described herein are available, OncoLoop can be readily applied to PCa in clinical practice.

Beyond PCa, OncoLoop is readily generalizable for predicting both cognate models and drug sensitivity for other cancers, as well as in non-cancer-related contexts. Notably, large-scale human, GEMM, and/or PDX cohorts are available for most cancer types. In addition, we have already generated genome-wide perturbational profiles in cell lines, comprising representatives for 23 distinct tumor subtypes (see (27) for partial coverage of the PanACEA database). Critically, the ability to stratify drug sensitivity in precisely identified molecular subtypes (pharmacotypes), if further validated, may lead to rapid design of basket and umbrella trials, including using adaptive design approaches to efficiently replace baskets that fail to validate.

Indeed, OncoLoop can support rapid-turnaround co-clinical studies to both identify and validate optimal drugs for a given patient in a wide variety of cancer contexts. In particular, once appropriate models and drug perturbation data are available, as should be achievable for many cancer types, the implementation of OncoLoop requires only transcriptomic data from a patient’s tumor or biopsy, which would first be matched to the preexisting models to identify those that are the highest fidelity (i.e., most significant MR-matched). Drugs predicted to be optimal inverters of the MR activity of a given human tumor and its cognate model (MR-inverter drugs) could then be validated in the preexisting models and, in the scenario in which FDA approved drugs are evaluated, adapted to clinical practice within a relatively short time-frame.

Nonetheless, there are also several caveats that will benefit from further refinements. First, despite the benefit of using FDA-approved drugs for rapid translation to clinical practice, the down-side of focusing only on drugs that are FDA-approved or in Phase II/III trials, is that such drugs may not be optimal for targeting the MR proteins most relevant for the tumors. Therefore, in future studies, extension to additional experimental agents may expand the repertoire of effective drugs, especially considering the new classes of PROTACS (54) and antisense agents (55). Additionally, drug perturbation profiles are limited by the availability of appropriate cell lines MR-matched to human patients. For example, the current study does not adequately inform on drugs that target NEPC tumors since neither the LNCaP nor the DU145 cell line recapitulate the MR-activity signature of this subtype. This can be addressed by generating drug perturbation profiles from primary, NEPC patient derived cells or organoids. Moreover, despite the encouraging results reported here, MR signature analysis and MR-inverter drug predictions are not 100% accurate, as is the case for most machine learning methods. We thus envisage that larger scale validation will be required before OncoLoop predictions may be confidently used in the design of co-clinical, basket, or umbrella clinical trials.

Another key caveat is the largely heterogeneous nature of most tumors, which present molecularly distinct subpopulations with potentially equally distinct drug sensitivity. Thus, when used to analyze bulk tumor profiles, OncoLoop may miss the opportunity to nominate drugs targeting the less represented subpopulations, thus selecting for drug-resistant ones and ultimately leading to relapse. A possible way to overcome this is to perform OncoLoop analyses at the single cell level, which should enable prioritization of drugs for all detectable subpopulations (56). A second approach is to perform analysis on the post-treatment minimal residual tumor mass, which is likely highly enriched for resistant subpopulations (57, 58). Additionally, while OncoLoop predictions are transcriptome-based, recent results show that additional omics modalities, such as a patient’s mutational profile and protein structure, among others, can be readily integrated to further refine MR protein identification and drug prediction (22).

Beyond cancer, MR-based predictions have been validated in disease as different as Parkinson’s (59), ALS (60), alcohol dependency (61), and Type II diabetes (62), suggesting direct OncoLoop applicability to such contexts, as long as appropriate drug-perturbation data were available. Indeed, while human translation of drugs validated in a GEMM or PDX context has been reasonably effective in cancer, mouse models have largely failed to recapitulate the effect of drugs on human disease. In this context, OncoLoop’s quantitative fidelity metrics may help to identify more appropriate mouse models for drug validation.

The current design of OncoLoop relies on MR-based predictions of model similarity and drug sensitivity. However, alternative approaches could also be tested using the same framework presented here. For instance, transcriptome-based approaches have shown promising results in predicting sensitivity of human patients in a clinical context (63), while neural network-based methods trained on multi-omics data have shown promising results in translating drug sensitivity assays from a training set of cell lines and mouse models to an independent test set of PDX models (64). Similarly, transcriptome-based approaches for assessing model fidelity have also been proposed (65).

Taken together, our data shows that OncoLoop may provide a valuable contribution to the emergent field of precision medicine, by complementing rather than supplanting existing approaches based on its ability to couple effective drug and high-fidelity model predictions.

## Materials and Methods

### A resource of genetically engineered mouse models (GEMM) of prostate cancer

All experiments using animals were performed according to protocols approved by the Institutional Animal Care and Use Committee (IACUC) at Columbia University Irving Medical Center (CUIMC). As described in Table S1A, the genetically engineered mouse models (GEMMs) used in this study utilize the *Nkx3.1^CreERT2^*^/+^ allele to activate an inducible Cre recombinase in the prostatic epithelium (35). The *Nkx3.1^CreERT2^*^/+^ allele was crossed with various other mouse alleles to achieve conditional deletion or conditional activation in the prostatic epithelium, or with the Hi-Myc transgene in which MYC expression is regulated by a constitutively active Probasin promoter (39). For lineage tracing to visualize tumors and metastases *ex vivo,* mice were further crossed with a conditionally-activatable fluorescent reporter allele (*Rosa-CAG-LSL-EYFP-WPRE*) (42). All mice were maintained on a predominantly C57BL/6 background; since the focus of our study is prostate cancer, only male mice were used. Several of these GEMMs have been described previously (30–34). All multi-allelic strains are available from the Jackson laboratory (Table S1A).

#### Phenotypic analysis

All studies were performed using littermates that were genotyped prior to tumor induction. Mice were induced to form tumors at 2-3 months of age by administration of tamoxifen (Sigma-Aldrich, St. Louis, MO) using 100 mg/kg (in corn oil) once daily for 4 consecutive days. Following tamoxifen-induction, mice were monitored three times weekly, and euthanized when their body condition score was <1.5, or when they experienced body weight loss ≥ 20% or signs of distress, such as difficulty breathing or bladder obstruction. Control (non-tumor induced) mice received only vehicle (i.e., corn oil) and were otherwise monitored in parallel with tumor-induced mice. As indicated, surgical castration was performed at 1-10 months after tumor induction (Table S1A). Where indicated, tumor volume was monitored by Magnetic Resonance Imaging (MRI) using a Bruker Biospec 9.4T Tesla Small Animal MR Imager located within the mouse barrier facility in conjunction with the Oncology Precision Therapeutics and Imaging Core (OPTIC) of the Herbert Irving Comprehensive Cancer Center (HICCC). Images were generated using ParaVision 6 software for preclinical MRI (Bruker) and reviewed by an MRI physicist. Volumetric analysis was done using 3DSlicer software (http://www.slicer.org).

At the time of sacrifice, tissues were collected and YFP-positive prostatic tumors and metastases were visualized by *ex vivo* fluorescence using an Olympus SZX16 microscope (Ex490–500/Em510–560 filter). For histopathological analysis, tissues were fixed in 10% formalin (Fisher Scientific, Fair Lawn, NJ), and hematoxylin and eosin (H&E) and immunostaining were done using 3 μm paraffin sections as described (30, 34). Antibodies used in these studies were: Androgen receptor (Abcam ab133273, 1:200; Cambridge, MA), Ki67 (eBioScience 14569882, 1:500; San Diego, CA), Pan-cytokeratin (Dako Z0622; 1:500; Santa Clara, CA) and Vimentin (Cell Signaling 5741S, 1:200; Danvers, MA). Images were captured using an Olympus VS120 whole-slide scanning microscope. Histopathological scoring of GEMM prostate cancer phenotypes was assessed independently by two pathologists (AR and SdB) based on evaluation of both the H&E-stained and adjacent immuno-stained tissues; these analyses are summarized in Table S1A,B.

#### Generation of allografts

Following protocols approved by IACUC at CUIMC, allografts were generated by transplanting freshly dissected prostate tissues from the GEMMs subcutaneously into the flank of male Athymic nude mice (Hsd:Athymic Nude-Foxn1nu, Envigo, Boyertown PA). Briefly, at the time of euthanasia, tumors were collected and cut into < 2mm pieces. Tissues were rinsed twice in cold sterile Dulbecco’s Modified Eagle Medium (DMEM; Thermo Fisher Scientific, Waltham, MA), soaked in Matrigel (Corning; Corning, NY), and implanted into the flanks of male hosts. Allografted tumors were harvested when the tumor size reached 2 cm or earlier if the body condition score of the host mouse was <1.5 or the mice exhibited signs of distress. At time of harvest, allografted tumors were passaged to new hosts and/or snap frozen in 80% DMEM, 10% fetal bovine serum (FBS, Gemini Bio-Products, West Sacramento, CA, USA), and 10% DMSO. Alternatively, the harvested allografts were fixed in 10% formalin for histopathological analyses, as above. A summary of allografted tumors is provided in Table S1A; as part of this GEMM resource, allografted tumors will be made available upon request by contacting the authors.

#### Generation of organoids

Mouse tumor organoids were generated as described (66). Briefly, freshly dissected prostate tissues were dissociated in DMEM containing 5% FBS and 1x collagenase/hyaluronidase (STEMCELL Technologies, Vancouver, BC, Canada), followed by trypsinization with 0.25% trypsin-EDTA (Gibco, Carlsbad, CA) at 4 °C for 1 h. Trypsinization was stopped by addition of Hanks’ Balanced Salt Solution (HBSS) media (STEMCELL Technologies) containing 2% FBS and the tissue was further dissociated by treatment with dispase (5 mg/mL)/DNaseI (0.1 mg/mL) solution (STEMCELL Technologies). The cell suspension was filtered using a 40 µm cell strainer (Falcon; Corning, NY, USA), resuspended in HBSS containing 2% FBS (Gemini Bio-Products, West Sacramento, CA) and subjected to fluorescence activated cell sorting (FACS) using a BD-FACS Aria cell sorter (BD Biosciences, Franklin Lakes, NJ) to collect the YFP-lineage marked prostate cells. Sorted cells were plated in low-attachment 96-well plates at 10,000 cells per well and cultured in organoid culture medium containing: hepatocyte medium (Corning) with 1x Glutamax (Gibco), 10 ng/mL EGF (Corning), 10 µM ROCK inhibitor (STEMCELL Technologies), 5% heat-inactivated charcoal-stripped FBS (Thermo Fisher Scientific), 100 nM DHT (Sigma-Aldrich), 5% Matrigel (Corning) and 1x antibiotic-antimycotic (Gibco). Organoids were harvested by centrifugation and embedded with Histogel (Thermo Fisher Scientific) (66), and processed for histopathological analyses, as above. A summary of organoids is provided in Table S1A. Although in the current study preclinical validation focused was done using the allograft models *in vivo*, as part of the GEMM resource, the mouse organoids will be made available upon request by contacting the authors.

### Transcriptomic analysis of GEMMs

RNA-seq data was generated from n = 136 GEMM-derived tumors (GEMM-DTs) and GEMM-derived normal prostate tissue. RNA was prepared from snap-frozen samples by homogenization in TRIzol Reagent (Invitrogen; Waltham, MA) using a Fisherbrand™ 150 Homogenizer (Fisher Scientific, Pittsburg, PA Catalog number 15-340-168) followed by extraction using the MagMAX-96 total RNA isolation kit (ThermoFisher Scientific, Waltham, MA, USA). Total RNA was enriched for mRNA using poly-A pull-down; only RNA samples having between >200 ng and 1μg and with an RNA integrity number (RIN) > 8 were used. Libraries were made using an Illumina TruSeq RNA prep-kit v2 or TruSeq Stranded mRNA library prep kit, and sequenced using an Illumina HiSeq2500/4000 or NovaSeq6000. RNA-seq profiles were generated by mapping mRNA reads to the mouse reference genome (version GRCm38 mm10), using *kallisto* v0.44.0 (67). Sequence-based bias correction was applied to pseudo-align the reads and quantify transcripts abundance in each sample. The resulting raw reads were mapped using the ENSEMBL gene model and summarized to transcript per million (TPM) to remove gene length bias thus allowing for inter and intra-sample comparison between genes. Raw and processed data were deposited in GEO (GSE186566).

#### Network analysis

A GEMM PCa-specific regulatory network (interactome) was reverse engineered from the resulting GEMM-DT and GEMM-DP RNA-seq profiles (n = 136) using ARACNe-AP (68), the most recent implementation of the ARACNe algorithm (29) (Table S2A). Default values for the algorithm were used for the number of bootstraps (n = 200), Mutual Information (MI) *P*-value threshold (P ≤ 10^-8^), and Data Processing Inequality (DPI) (enabled = yes). A total of *n* = 3,163 regulatory proteins were selected from Gene Ontology (GO)-generated protein sets, including n = 1,046 Transcription Factors (TF)—“DNA-binding transcription factor activity” (GO:0003700)—and *n* = 2,217 co-Transcription Factors and chromatin remodeling enzymes—“Transcription coregulator activity” (GO:0003712), “DNA binding” (GO:0003677), “Transcription factor binding” (GO:0008134), “Regulation of transcription, DNA-templated” (GO:0006355), “Histone binding” (GO:0042393), and “Chromatin organization” (GO:0000790)—see (69, 70).

#### GEMM-DT analyses

For all subsequent analyses, we selected a subset of n = 91 GEMM-DTs that most faithfully recapitulated pathophysiologically-relevant prostate cancer phenotypes, based on their histopathology (Table S1B).

#### Interactome comparison

The expression matrix of the GEMMS was converted to gene expression signatures using Median Absolute Deviation (MAD) and independently analyzed with VIPER using both a prior GEMM-specific interactome (20), generated from Illumina gene expression microarrays data, *I*_Old_, and the new interactome described above, *I*_New_. For each GEMM-DT sample (n = 91) the 25 most activated (25↑) and 25 most inactivated (25↓) MRs were identified. The mean of the absolute Normalized Enrichment Score (NES) across all 50 MRs was used to assess the interactome’s ability to identify sample-specific, statistically significantly differentially active proteins. The NES probability densities and expected values were also computed and plotted (Fig. S4A), showing that the new interactome is more effective in identifying differentially active proteins compared to the prior one (*P* < 2.2×10^-16^, as assessed by a two-sample Kolmogorov-Smirnov test).

#### GEMM-DT subtype identification

We used the VIPER algorithm (23) to transform the RNA-seq profile of each GEMM-DT into differential protein activity profiles, using regulatory protein targets (regulons) captured by the GEMM interactome. To avoid bias due to different regulon sizes, regulons were pruned to include only the 50 highest likelihood targets, as recommended in (23), and regulons with < 50 targets were excluded from the analysis. From 2,858 regulatory protein regulons recovered by ARACNe inference, only 64 were removed due to inadequate regulon size. The resulting regulatory network comprising regulons of 2,794 regulatory proteins was used for subsequent GEMM-related analysis.

Since this analysis was aimed at stratifying disease subtypes by identifying the proteins that are most differentially active between subtypes, we generated sample-specific gene expression signatures (GES) based on the *z*-score of differentially expressed genes in each GEMM-DT vs. the average of all GEMM-DTs as the reference distribution. This approach is designed to maximize intra-cohort differences. Results are summarized in Table S2B.

We performed Uniform Manifold Approximation and Projection (UMAP) (71) for dimensionality reduction analysis on the 91 GEMM-DT protein activity profiles, after selecting the most statistically significant principal components (PCs), by applying the JackStraw algorithm (72). Six statistically significant PCs were identified (*P* ≤ 0.01) and used for downstream analysis. Next, we built a Shared Nearest Neighbor (SNN) Graph (73) using the Seurat v3.1.2 (74) implementation of the algorithm. For the SSN algorithm we used *k* = 7 as number of neighbors for graph construction. Finally, cluster analysis was performed using the Louvain algorithm (75). To select the optimal *resolution* parameter for the Louvain algorithm, we ran a grid search across the range [0.01, 2] in steps of 0.05 and used the average Silhouette Score (76) and co-segregation of clinical parameters (e.g., metastatic samples, NEPC samples, etc.) across the possible cluster solutions as cost function to optimize the number of clusters. The resulting data are provided in Table S2C. The optimal 5-cluster solution was found at a resolution *ρ* = 0.41. Each cluster should thus represent a molecularly distinct GEMM-DT subtype.

To identify the proteins that were most differentially active in each subtype—*i.e.*, representing candidate, subtype-specific Master Regulator (MR) proteins—we integrated the z-scores representing the VIPER-measured differential activity of each protein across all subtype samples having silhouette score > 0.25, using Stouffer’s method. To avoid clutter, we visualized the 10 most activated proteins for each subtype (P ≤ 0.01, Bonferroni Corrected (BC)) as representative candidate positive MRs (Fig 3A and Table S2C).

#### Pathway analysis

Gene sets representing canonical pathways and Cancer Hallmarks were selected from the Broad MSigDB (**47**), KEGG (**48**), and REACTOME databases (**49**), see http://www.gsea-msigdb.org/gsea/msigdb/collections.jsp. To convert murine genes to human genes, for pathway analysis, gene identifiers were mapped to their human-mouse orthologs using the R biomaRt services (**77**). Gene Set Enrichment Analysis (GSEA) was then computed using the analytical Enrichment Analysis Algorithm (aREA) function from the R VIPER package (**23**) and used to assess the enrichment of each pathway or Cancer Hallmark gene set in genes encoding for proteins that were differentially active in each cluster. Gene sets with < 20 or > 250 genes were excluded as statistically underpowered or unspecific, respectively. Enrichment results are shown in Table S2D.

### Patient-Derived xenograft (PDX) model analysis

Following protocols approved by IACUC at CUIMC, LuCaP PDX lines 73, 77, 78, 81 and 147 (15) were continuously maintained by passage in male CIEA NOG mice (Taconic, Germantown, New York, USA). Xenografts were harvested when the tumor size reached 2 cm or earlier if the body condition score of the host mice were <1.5 or if they exhibited signs of distress. When the Xenograft tumors were 7-8mm in diameter, the host mice were castrated or left intact (mock surgery). To generate additional variability for effective interactome assembly, three days later, the mice were treated with either vehicle or one of 13 selected perturbagens described in (20), once daily, for 5 days. On the afternoon of the fifth day of treatment, mice were euthanized and tumors were collected and snap-frozen in liquid nitrogen, for a total of 140 samples (5 models, 14 treatments, 2 castration states).

RNA extraction and library preparation were performed as described for the GEMM-DT cohort and libraries were sequenced on the Illumina HiSeq2000 instrument. We used RTA (Illumina) for base calling and CASAVA (version 1.8.2) for converting BCL to fastq format, coupled with adaptor trimming. Reads were mapped to a reference genome (Human: NCBI/build37.2) using Tophat (version 2.0.4) (78) with 4 mismatches (--read-mismatches = 4) and 10 maximum multiple hits (--max-multihits = 10). The relative abundance (aka expression level) of each gene and splice isoform was estimated using cufflinks (version 2.0.4) (79) with default settings. Expression profiles for both the raw and processed data have been deposited in GEO (GSE184427). A Xenograft interactome was generated from the 120 highest quality xenograft-derived gene expression profiles using ARACNe-AP (68), as described for the GEMM cohort. Results are provided in Table S7A.

### Human patient cohort analysis

For the analysis of human tissue from PCa tumors and normal prostate, we collected n = 790 RNA-seq profiles from published sources, including profiles from: (*a*) 245 normal prostate tissues from the GTEx consortium (version 8, date 2017-06-05) (50); (*b*) 333 treatment-naïve, clinically-annotated primary prostate adenocarcinoma samples in The Cancer Genome Atlas (TCGA) PCa cohort (26); and (*c*) 212 metastatic biopsies from heavily pretreated mCRPC patients from the Stand Up to Cancer cohort (SU2C) (25). In addition, we refer to a cohort of 49 metastatic biopsies from patients with castration-resistant adenocarcinoma (adeno-CRPC) or neuroendocrine prostate cancer (NEPC) from which a NEPC signature was derived as discussed in (11).

#### TCGA interactome

To generate the primary prostate tumor interactome we used data from the full set of 498 patients in the TCGA PCa cohort (26). RNA-Seq counts data were downloaded from https://portal.gdc.cancer.gov/repository using TCGAbiolinks for R (80). Regulatory network analysis was performed as described for the GEMM interactome, using a total of n = 2,554 regulatory proteins were selected into manually curated protein sets, including n = 1877 Transcription Factors (TF) (GO:0003700) and *n* = 677 co-Transcription Factors or chromatin remodeling enzymes (GO:0003712), see (69, 70) Data are provided in Table S3A. The interactome was then generated as described for the GEMM-DT samples.

#### SU2C interactome

RNA-seq profiles of this cohort were collected on October 4, 2019 from cBioPortal as FPKM measurements, as released in the 2019 SU2C-mCRPC dataset (25). We restricted our analysis to n = 212 patients for which RNA-seq data had been generated using the Agilent SureSelect Human All Exon V4 reagents. FPKM were transformed to TPM and regulatory network generation was performed as described above. Data are provided in Table S3B.

#### Generation of differential gene expression signatures

To identify gene expression signatures of tumorigenesis for protein activity analysis—including for MR-based patient/GEMM-DT matching and to identify MR-inverter drugs—tumor samples from each cohort (*n* = 333 for TCGA and n = 212 for SU2C), was compared to the same *reference control* dataset, comprising RNA-seq profiles of 245 normal human prostates from the GTEx cohort, as above, and 26 normal murine prostates, following mouse-to-human gene conversion. The latter were obtained from wild-type and *Nkx3.1* heterozygous and homozygous null mice (Table S1B). To allow sequencing depth-independent comparison of tumor vs. *control* samples, including for low-depth PLATE-seq profiles (∼1M reads), we normalized each sample to an equivalent library size of 10^6^ reads, after filtering the data to retain only genes represented across all analyzed datasets, followed by Log transformation. We used the R *limma* package (81) to fit a linear model to each gene using the *control* dataset as reference. Next, for each gene, we computed *P*-values and moderated t-statistics using the empirical Bayes method with parameter *robust* set to TRUE. For each sample, we used a vector of these statistics to generate a z-score vector representing the differential expression of each gene in a tumor sample versus *reference control* (signature). The GEMMs cohort was processed identically, after cross-species conversion of gene identifiers, which was done as described in the Pathway analysis section above.

#### Generation of protein activity profiles

Differential protein activity was measured by VIPER analysis of differential gene expression signatures—i.e., each tumor vs. the average of the normal samples control reference—using the corresponding cohort-specific regulatory network. All cohorts were processed using the same parameters. Protein activity profiles are provided in Tables S3C,D.

### OncoMatch analysis

To assess the fidelity of an individual GEMM, cell line or PDX (i.e., tumor models) to an individual human tumor sample, we used their protein activity signatures to compute the NES of the 25↑+25↓ most differentially active proteins (*MR-signature*, hereafter), as assessed by VIPER analysis of a human sample, in proteins differentially active in the model sample, using the aREA algorithm, as shown in (24). A fixed MR number was selected to avoid bias due to set size in assessing the NES, while the specific number (*i.e.*, n = 50) was selected based on the results of a recent pancancer study (22) where an average of 50 MR was found to be sufficient to canalize the effect of the functionally relevant genetic alterations, on a sample-by-sample basis, across 20 TCGA cohorts. Murine proteins were *humanized* (*i.e.*, mapped to human proteins), as described in the Pathway analysis section, via custom R functions to query the BiomaRt servers (77). The aREA algorithm (23) was then used to estimate Normalized Enrichment Scores (NES) that were finally converted to *P*-values. The conservative Bonferroni method was used to correct for multiple hypothesis testing and the value *S_F_* = −Log_10_ *P* was used as an MR conservation-based fidelity score. Tumor models with fidelity score *S_F_* ≥ 5 (i.e., *P* ≤ 10^-5^) were then selected as high-fidelity cognate models.

For drug validation purposes, the GEMM-DTs with the highest, statistically significant fidelity score (P ≤ 10^-5^) were used as optimal models for patient-relevant drug validation. All available GEMM-DTs (n = 91) were tested as potential high-fidelity models of all TCGA and SU2C cohort samples. Results are provided in Tables S4A and S4B, respectively, and in Figure 4, where we show both the top 5 highest fidelity models per patient, as well as all additional statistically significant models (P ≤ 10^-5^).

### Human PCa cancer cell line selection

To generate drug perturbational profiles, we first identified cell lines representing the highest fidelity models of TCGA PCa samples. Cell Line RNA-seq profiles were downloaded from the Broad Institute web portal (https://data.broadinstitute.org/ccle/) from the Cancer Cell Line Encyclopedia (CCLE) as transcript per million (TPM) measurements with timestamp version 20180929. From these data, which comprise the RNA-seq profiles of 1,019 cell lines, we generated gene expression signature (GES) representing the z-score of genes differentially expressed in each of 8 PCa cell lines in CCLE against the average of all CCLE cell lines as a control reference. Differential gene expression signatures were then VIPER transformed into differential protein activity profiles using the metaVIPER algorithm (82) to select either the TCGA or the SU2C interactome as the optimal human PCa network on a protein-by-protein basis.

For androgen-dependent tumors, LNCaP emerged as an optimal model (P ≤ 10^-5^) for >70% of the indolent TCGA samples. The DU145 cell line, while matching a much smaller fraction of the TCGA samples was identified as an optimal match (P ≤ 10^-5^) for TCGA samples presenting high Gleason Score, low AR activity, and high CENPF and FOXM1 activity—two proteins previously reported as synergistic determinants of malignant PCa (20), which were also aberrantly activated in this cell line. These cell lines were thus selected for drug perturbation assays.

#### Dose-response curve (DRC) generation

To determine the maximum sublethal concentration of each drug—defined as its 48h EC_20_ concentration—cells were seeded onto 384-well tissue culture plates (Greiner 781080 Monroe, NC; columns 1-24, 50 μL volume) at a density of 1,000 cells per well. Plates were incubated at room temperature for 30 minutes, and then overnight in an incubator (37 °C, 5% CO_2_). The next morning (approximately 12 hours after seeding) compounds were added using the Echo 550 acoustic dispenser. After 40 μL of media were removed from each well, each drug was dispensed in ten doses, with each plate run in triplicate. After 48hrs, 25 μL of CellTiter-Glo (Promega Corp, Madison, WI) were added to each well. Plates were shaken for 5 minutes before enhanced luminescence readout (EnVision, Perkin Elmer, Shelton, CT). All values were normalized by internal controls contained on each plate and analyzed for EC_20_ determination using Pipeline Pilot, Dassault Systems.

#### Drug perturbation analyses

Compared to DRC generation, more cells were needed to isolate sufficient total RNA-seq profile generation. For this purpose, cells were thus seeded onto 384-well tissue culture plates, as above, at a density of 4,000 cells per well. The next morning (approximately 12h later), compounds were added using the Echo 550 system as above. Each drug was dispensed at the smaller of its previously determined EC_20_ or its calculated C_max_ (maximum tolerated serum concentration), but never above 10 μM. The C_max_ was used to avoid perturbing cells with non-physiologically relevant drug concentrations. After 24 hours, plates were spun down at 295 × g for 1 minute. Media was removed and cells were re-suspended in 30 μL of Turbo Capture Lysis (TCL) buffer (Qiagen) containing 1% beta-mercaptoethanol (BME). Finally, PLATE-Seq analyses was performed as described (51) at 24h after drug perturbation. Cells were harvested at 24h after perturbation with 335 compounds, including 117 FDA-approved oncology drugs and 218 experimental compounds (Table S5A). The screen was performed by the Columbia JP Sulzberger Genome Center High-Throughput Screening facility of the Herbert Irving Comprehensive Cancer Center (HICCC), which makes PLATE-seq screens available to the entire research community.

While the study used only profiles from DU145 cells—since they were a better model for the aggressive mCRPC samples in the SU2C cohort—drug perturbation profiles from both LNCaP and DU145 cells will be accessible from GEO199800.

#### Data analysis

PLATE-Seq profiles were mapped to the human reference genome version Grch38 using STAR aligner (83). Variance stabilization, from the DESeq2 package for R (84), was used to normalize the data from each plate. To correct for potential cross-plate batch effects, we used the function *combat* from the sva package for R (85) to compute the batch-corrected normalized gene expression and to fit a linear model for each compound against six plate-matched DMSO controls, included to minimize the need for cross-plate normalization. We used the *limma* package for R (81) to fit the linear model and to compute *P*-values and moderated t-statistics for each gene. For each compound, we used a vector of these statistics to generate a gene expression profile of z-scores representing the compound effect as differential between post-and pre-treatment. To account for tumor context-specificity, we performed VIPER-inference on the DU145 screening data using the SU2C interactome (as described above). The drug perturbation data are provided in Table S5B.

### OncoTreat Analysis

OncoTreat analysis was performed as described in (21). Briefly, drug-mediated MR inversion was independently assessed for each patient in the SU2C cohort and each GEMM as follows: for each sample and for each drug, we assessed the NES of the sample’s MR-activity signature (25↑+25↓ most differentially active proteins) in proteins differentially inactivated and activated in drug-treated vs. vehicle control treated cells, respectively, using the aREA algorithm (23). NES values were then converted to P-values, and Bonferroni corrected to account for multiple hypothesis testing. An efficacy score was computed *S_E_* = −Log_10_ *P* and used to identify MR-inverter drugs (*S_E_* ≥ 5, corresponding to *P* ≤ 10^-5^). Results of OncoTreat analyses for the SU2C and GEMM cohorts are provided in Table S5C and Table S5D, respectively.

### OncoLoop Analysis

The OncoLoop algorithm leverages a tripartite graph *TPG* with nodes representing patients (*P_i_*), GEMM-DT (*G_j_*) and drugs (*D_k_*), respectively, and edges represent statistically significant GEMM-patient fidelity score *S_M_* (*P_i_*, *G_j_*) ≥ 5, GEMM-drug MR-inverter score *S_E_* (*G_j_*, *D_k_*) ≥ 5 and patient-drug MR-inverter score *S_E_*(*P_i_*, *D_k_*) ≥ 5. All closed 3-node loops including a patient, a GEMM-DT, and a drug are considered as statistically significant PGD-Loops. These were then ranked based on the Stouffer’s integration of the z-scores corresponding to the *S_M_* (*P_i_*, *G_j_*), *S_E_*(*G_j_*, *D_k_*), and *S_E_*(*P_i_*, *D_k_*) values of the loop, which can then be converted back to a P-value.

Analysis of 212 human samples, 91 GEMM samples, and 335 drug perturbations yielded 668,138 statistically significant PGD-Loops, which are summarized in Table S6. For each patient, the optimal GEMM-DT for validation and most likely MR-inverter drug(s) for both patient and GEMM-DT can be identified by selecting among all the PGD-Loops that include that patient, those with the highest Stouffer’s integrated rank.

### Preclinical Validation of OncoLoop-predicted Drugs

#### Candidate drug selection

To select candidate drugs for *in vivo* validation we analyzed the 668,138 significant PGD-Loops produced by OncoLoop analysis of SU2C cohort patients, as described in the previous section. Specifically, to maximize the translational potential of selected drugs, we focused on loops restricted to (*a*) including only FDA-approved drugs; (*b*) including drugs tested at a physiologically relevant concentration ≤ 1 *μM*, and (*c*) including drugs predicted as MR-inverters for ≥ 50% of SU2C samples (n=212), thus suggesting broad applicability. The analysis yielded 16 drugs, which were then sorted by patient coverage—i.e., the fraction of patients in PGD-Loops comprising the specific drug (Figure 6A,B).

#### Candidate drug validation

For validation of candidate drugs, we tested their ability to inhibit tumor growth *in vivo* in cognate GEMMs. Cognate GEMMs for validation were selected based on their being (1) included in the closed loops (i.e., matching both the drug and the SU2C patient) and (2) readily available as allograft models. Based on these criteria, we selected two allograft models: *NPp53^mut^* CMZ315 and *NPM* CMZ150 (Table S1B). In addition, we validated drug candidates in the MR-matched human PDX LUCAP-73 model (Table S7B) (15). For the allograft tumors, 20-40 mg pieces from passage 2 tumors were grown in the flanks of male NCr *nude* mice (Envigo), as above. For the LUCAP-73 xenografts, 100 mg pieces were grown in R2G2^®^ mice (B6;129-Rag2tm1FwaII2rgtm1Rsky/DwlHsd, Envigo). Tumors were monitored by caliper measurement twice weekly and tumor volumes were calculated using the formula [Volume = (width)^2^ x length/2]. When tumors reached 100-200mm^3^, mice with similar mean tumor volume were randomized into vehicle and treatment groups.

Pharmaceutical grade compounds for the 4 candidate drugs, namely Temsirolimus (S1044), Trametinib (S2673), Panobinostat (S1030), Bortezomib (S1013), and Cabazitaxel (S3022) as a control for standard of care, were purchased from Sellekchem (Houston, TX). Stock solutions were made by dissolving the compounds in 100% ethanol (Temsirolimus) or DMSO (all others) that were frozen as single-use aliquots at −80°C. Stock solutions of the drugs were freshly diluted before use in vehicle (5%PEG400, 5%Tween80 in PBS). For each candidate drug, the dosage, mode of delivery and schedule was chosen based on previous reports and were as follows: Temsirolimus (20mg/kg) (86), Trametinib (1mg/kg) (87), Panobinostat (15mg/kg) (88), Bortezomib (1mg/kg) (89), and Cabazitaxel (10mg/kg) (90) Drugs were administered via intraperitoneal delivery (i.p.) 3 times/week in non-consecutive days (Temsirolimus, Panobinostat, Bortezomib, and Cabazitaxel), or by oral gavage 5 times/week in consecutive days (Trametinib). Tumors were harvested when the tumor size of vehicle treated mice reached 2 cm or earlier if the body condition score of the host mice were <1.5 or if they exhibited signs of distress. Growth curves were analyzed using two-way ANOVA with Dunnet’s multiple comparisons test, compared to vehicle-treated controls. Mice with tumors greater than 2.0 cm, or having 20% weight loss from baseline, or with body condition scoring of 1.5 or less were euthanized by CO2 inhalation followed by cervical dislocation to verify death. Tumors were fixed in 10% formalin to be processed for histology, or snap-frozen in liquid nitrogen.

#### Pharmacokinetic analyses

To quantify the abundance of drug in the treated tumors, pharmacokinetic studies were performed using Ultra Performance Liquid Chromatography-Tandem Mass Spectrometry (UPLC-MSMS) by the Biomarkers Core Laboratory at CUIMC. Drugs were extracted from 10 mg snap-frozen tissues from tissue homogenates spiked with internal standards by liquid-liquid extraction using diethyl ether: ethyl acetate (30:70) and resuspended in 20% methanol for LCMS analysis. LCMS analysis was carried out on a Waters Xevo TQ MS integrated with ACQUITY UPLC system (Waters, Milford, MA, USA). The system was controlled by MassLynx Software 4.1. Single assays were used for bortezomib and Temsirolimus whereas Panobinostat and Trametinib were measured in two separate LCMS reactions. The lower limit of quantitation was 2.5ng/mL (Trametinib, Borteozomib) and 5ng/mL (Temsirolimus, panobinostat). The intra-assay precision was as follows: Temsirolimus 5.34%, panobinostat 5.06%, Trametinib 4.83% and bortezomib, 3.20%.

#### Pharmacodynamic analyses

Pharmacodynamic assays were performed at an early time point following *in vivo* drug treatment to assess each drug’s ability to recapitulate the computationally predicted MR inversion. For this purpose, allografted tumors were grown in host mice, as above. When the tumors reached the size of 100-200mm^3^, the host mice were treated with drug, as above, once daily for three consecutive days at the above-indicated doses. On the morning of day 4, tumors were collected and snap-frozen for RNA extraction or fixed in formalin. RNA sequencing was performed as described above for GEMMs. The data is deposited in GEO (GSE186566).

RNA-seq profiles were pseudo-aligned to mouse reference genome version GRCm38 mm10, using *kallisto* v0.44.0 (67) with sequence-based bias correction. Transcript abundance was quantified and summarized in gene counts mapped to the ENTREZ gene model. We used limma *voom* to transform count data to log2-counts per million (log_2_CPM) and account for mean-variance relationship (81). Next, we fitted gene-wise linear models and computed moderated t-statistics using empirical Bayes. Contrasts were created between treated cells and cell treated with vehicle only. Sample biological replicates (n=3) were used as blocking factor in the experimental design. To generate a gene expression signature (GES) of treatment-induced transcriptional changes we mapped the differential gene expression data (i.e., FDR-adjusted p-values and log-Fold Change scores) to a normal distribution. Analyses of treatment-induced MR-reversal was performed as described above.

### Statistical analyses

Statistical analysis was performed using GraphPad Prism software (Version 9.3.1) and R-studio (0.99.902, R v4.0.2). Kaplan-Meier survival analysis was performed using two-tailed log-rank test compared to the NP model. Comparison of frequencies was done using two-tailed Fisher’s exact test or as described in figure legends. Subcutaneous tumor growth curves were analyzed using One-way ANOVA at the last time point compared to Vehicle treated tumors, adjusted for multiple comparisons with Dunnett’s test. All bars show the mean and error bars SD. GSEA were computed by custom code implemented as per (91) algorithm. *P-*values estimated using a null distribution of 1,000 permutations of random fixed-size gene sets; NES, normalized enrichment score. No statistical method was used to predetermine the sample size used for *in vivo* experiments.

### Data Accession

The following datasets are deposited in GEO: (*a*) The mouse gene expression profiles (raw and normalized data) (GSE186566) (*b*) The human PDX gene expression profiles (raw and normalized data) (GSE184427), and (*c*) The PLATE-seq data for the drug perturbation profiles in both LNCaP and DU145 cells (GEO199800).

## Author Contributions

**A.V.** Conceptualized, designed and performed the computational analyses, and wrote manuscript.

**M.Z.** Conceptualized, designed and performed the generation and analysis of the GEMMs.

**J.M.A.** Conceptualized, designed and performed the drug validation studies, and wrote manuscript.

**F.N.A.** Designed and executed experiments, and performed analysis of the GEMMs.

**E.F.D., C.K., R.R., and S.P.** Designed and executed experiments, and performed data analysis related to the drug perturbation studies.

**M.S., A.M., and C.W.C.** Designed and executed experiments, and performed analysis of the PDX interactome.

**A.R.C. and S.d.B.** Performed histopathological analysis.

**J.Y.K.** Performed experiments.

**E.C.** Provided the PDX models

**M.J.A.** Performed data analysis.

**M.M.S.** and **M.A.R.** Supervisory role and data analysis.

**A.C.** Conceived the study, designed experiments, performed data analysis, and wrote manuscript.

**C.A.S.** Conceived the study, designed experiments, performed data analysis, and wrote manuscript.

## Acknowledgements

We thank Alvaro Aytes, Filippo Giancotti, Kenneth Olive, and Anil Rustgi, for comments on the manuscript. We are grateful to Sarah Bergen for assistance with PDX models, and Stephanie Afari for assistance with the GEMMs. Figure 1 was created with BioRender.com using an institutional license sponsored by Columbia University’s VP&S Office for Research.

These studies were supported by Flow Cytometry, Genomics and High Throughput Screening, and Oncology Precision Therapeutics and Imaging Core facilities which are funded in part through Herbert Irving Comprehensive Cancer Center support Grant P30 CA013696, the Biomarkers Core Laboratory at the Irving Institute for Clinical and Translational Research, home to Columbia University’s Clinical and Translational Science Award UL1TR001873, the Translational Research Unit, Institute of Pathology, University of Bern, and the PDX core at the University of Washington, which is supported by the PNW Prostate Cancer SPORE P50CA097186 and P01CA163227. Funding to support the deposition of mouse strains at The Jackson Laboratories was provided by the TJ Martell Foundation for Leukemia, Cancer and AIDS Research and the Prostate Cancer Foundation

This work was supported by the NCI Cancer Target Discovery and Development Program (U01 CA217858, to AC), the Cancer Systems Biology Consortium (U54 CA209997, to AC), NIH Shared Instrumentation Grants (S10 OD012351 and S10 OD021764 to AC), R01 CA173481 (to CAS), R01 CA183929 (to CAS), P01 CA265768R01 (to MMS), CA238005 (to MMS), U01CA261822 (to MMS), and a Prostate Cancer Challenge Award (to MMS and AC). AV was supported by a U.S. Department of Defense Early Investigator Research Award (W81XWH19-1-0337) and an Early Career Development Pilot Award NIH/NCI Cancer Center, funded through the Cancer Center Support grant, P30CA013696. MZ was supported in part by the National Center for Advancing Translational Sciences, National Institutes of Health, Grant Number UL1TR001873. JMA was supported by the Dean’s Precision Medicine Research Fellowship from the Irving Institute for Clinical and Translational Research at Columbia University Irving Medical Center (UL1TR001873), and a Prostate Cancer Foundation Young Investigator Award. MS was supported by NIH K99/R00CA194287. AM was supported by a Prostate Cancer Foundation (PCF) Young Investigator award. C.A.S. is an American Cancer Society Research Professor supported in part by a generous gift from the F.M. Kirby Foundation.

