## Supplementary Materials for "OncoLoop: A network-based precision cancer medicine framework"

Vasciaveo et al.

**Supplementary Figures**

**Figure S1:** Genomic alterations in prostate cancer represented in the GEMMs (related to Fig. 2)

**Figure S2:** Additional phenotypic analyses of the GEMMs (related to Fig. 2).

**Figure S3:** Phenotypic analysis of allograft and organoid models (related to Fig. 2).

**Figure S4:** Additional transcriptomic analyses of the GEMMs (related to Fig. 3).

**Figure S5:** Regulatory sub-networks of the GEMM clusters (related to Fig. 3).

**Figure S6:** Drug perturbation protein activity profiles from DU145 cells (related to Figs. 5, 6, 7).

**Figure S7:** Additional validation of drug candidates (related to Figs 6, 7)

**Supplementary Tables (provided separately)**

**Supplementary Table 1**: Phenotypic analysis of the GEMMs

A. Summary of GEMMs used in this study
B. Detailed description of individual mice analyzed by RNA sequencing and histopathology

**Supplementary Table 2**: Transcriptomic analyses of the GEMMs

A. GEMMs Interactome
B. Protein activity
C. Cluster analyses
D. Pathway analyses

**Supplementary Table 3**: Transcriptomic analyses of the human patient samples

A. TCGA Interactome
B. SU2C Interactome
C. Protein activity TCGA
D. Protein Activity SU2C

**Supplementary Table 4**: OncoMatch

A. OncoMatch TCGA

B. OncoMatch SU2C

**Supplementary Table 5.** Drug perturbation and OncoTreat analysis

A. Summary of drugs and concentrations used
B. Drug perturbation data for DU145
C. OncoTreat for drugs to SU2C patients using DU145
D. OncoTreat for drugs to GEMMs using DU145

**Supplementary Table 6.** OncoLoop summary table

**Supplementary Table 7.** Transcriptomic analyses of the PDX models

A. PDX interactome
B. OncoTreat for drugs to PDX using DU145 cell perturbation data

**Figure S1**

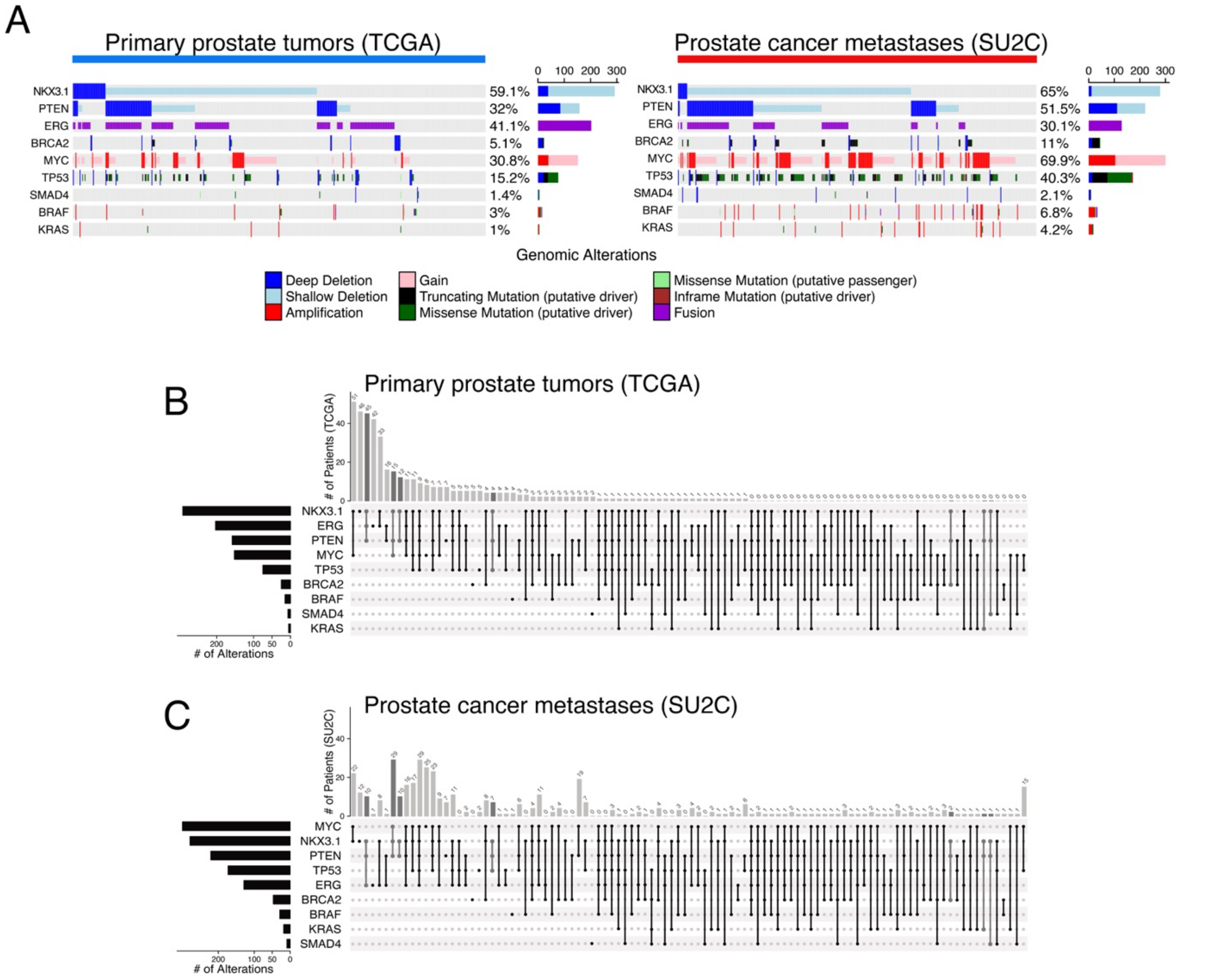


**Figure S1: Genomic alterations in prostate cancer as represented in the GEMM cohort (related to Figure 2)**

**A.** Oncoprint depiction showing the percentage of patients having genomic alterations that are represented in our GEMM cohort as depicted for primary prostate adenocarcinoma (TCGA, left pane) and prostate cancer metastasis (SU2C, right panel). **B,C.** Shown are the same alterations (rows) grouped by GEMMs (dark gray vertically connected dots) across patients that bear them in combination (columns) in the TCGA (**B**) and in SU2C (**C**) cohorts.

**Figure S2**

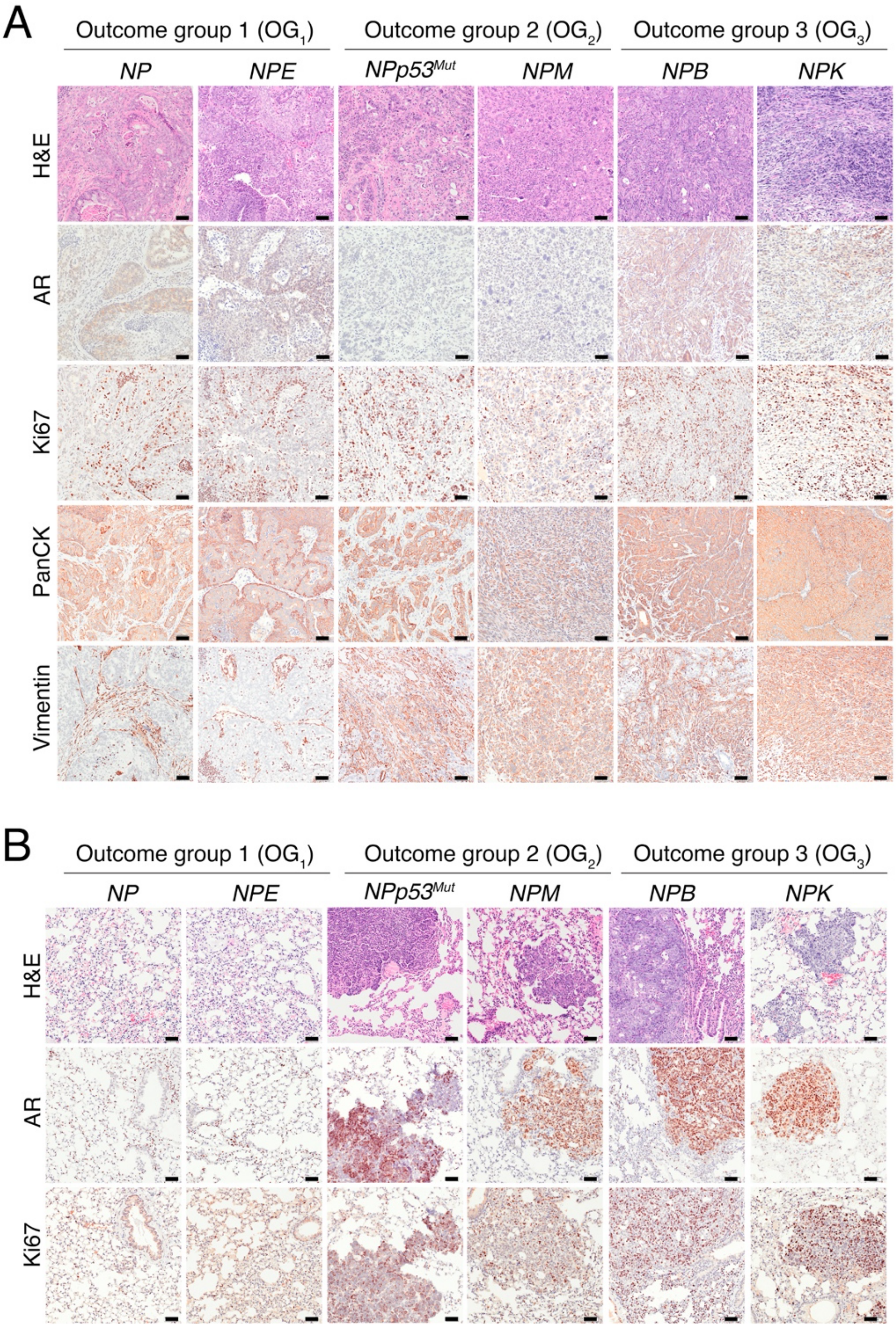
A
Outcome group 1 (OG1)
Outcome group 2 (OG2)
Outcome group 3 (OG3)
NP
NPE
NPp53Mut
NPM
NPB
NPK
H&E
AR
Ki67
PanCK
Vimentin
B
Outcome group 1 (OG1)
Outcome group 2 (OG2)
Outcome group 3 (OG3)
NP
NPE
NPp53Mut
NPM
NPB
NPK
H&E
AR
Ki67

**Figure S2: Additional phenotypic analyses of the GEMMs (related to Figure 2).**

**A.** Castration phenotype: Primary tumors from castrated mice of the indicated genotypes. **B.** Metastasis phenotype: Lung tissues from the indicated GEMMs with ($OG_2$, $OG_3$) or without ($OG_1$) metastases. Panels A and B show hematoxylin and eosin (H&E) and immunohistochemical staining of the indicated markers. Shown are representative images from analysis of ≥ 3 mice; scale bars represent 50µm.

**Figure S3**

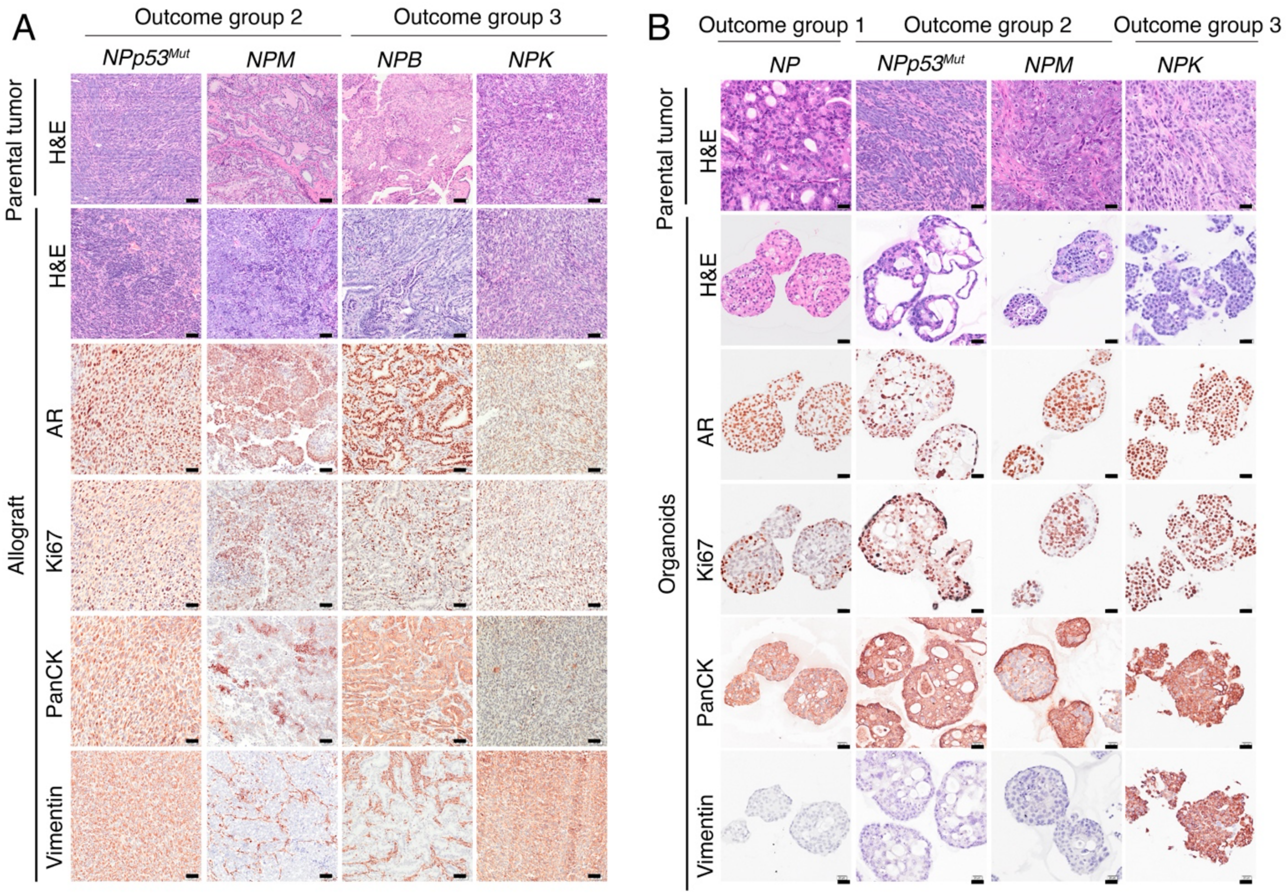


**Figure S3: Phenotypic analyses of allograft and organoid models derived from the GEMM cohort (related to Figure 2).**

**A,B.** Representative images for hematoxylin and eosin (H&E) and immunohistochemical staining of the indicated markers in allografts (**A**) and organoids (**B**) derived from the indicated GEMMs. All images represent the analysis of ≥ 3 cases; scale bars represent 50µm in **A**) and 20µm in **B**).

**Figure S4**

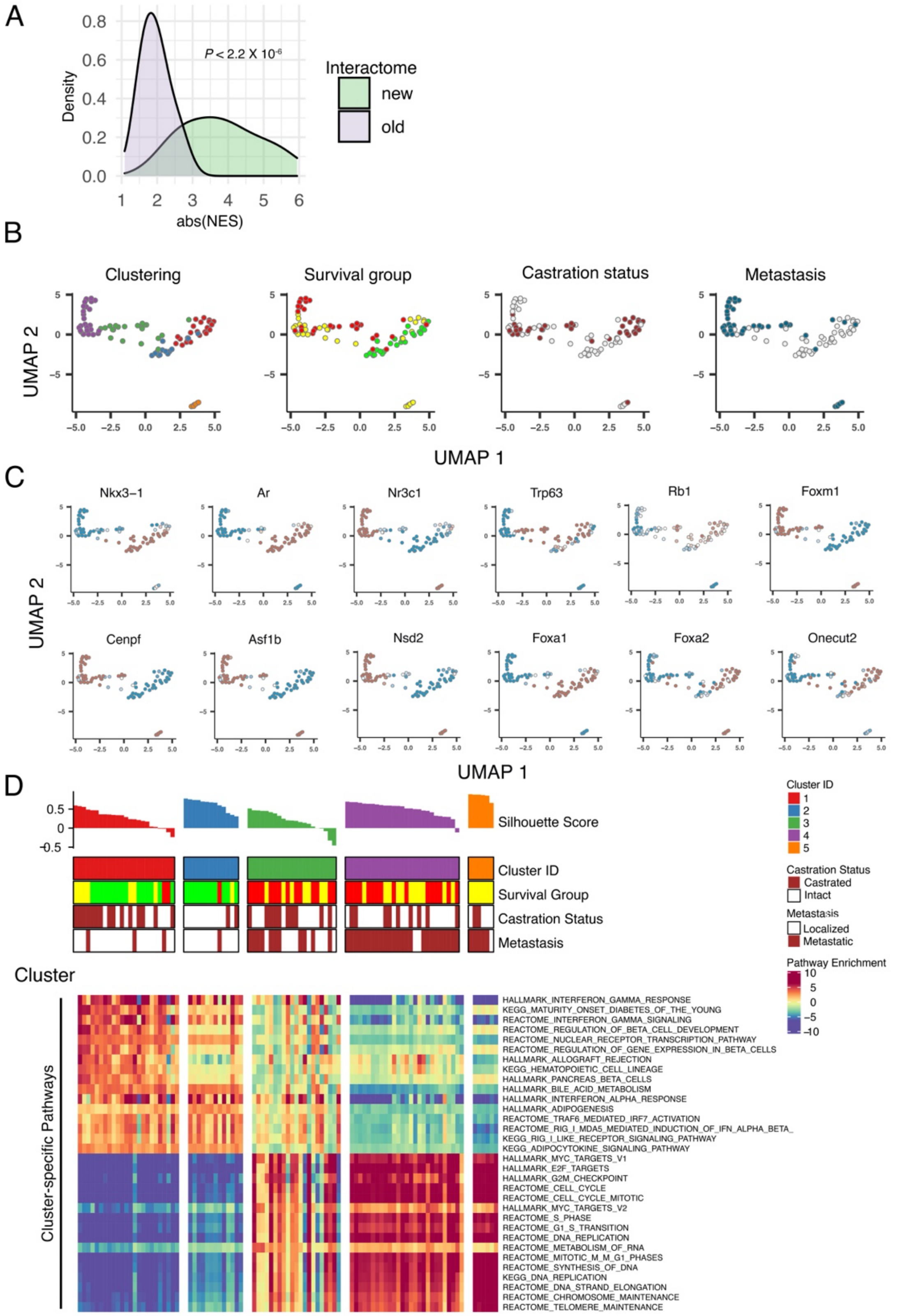
A
Density
abs(NES)
P < 2.2 X 10-6
Interactome
new
old
B
Clustering
Survival group
Castration status
Metastasis
UMAP 2
UMAP 1
C
Nkx3−1
Ar
Nr3c1
Trp63
Rb1
Foxm1
Cenpf
Asf1b
Nsd2
Foxa1
Foxa2
Onecut2
UMAP 2
UMAP 1
D
Silhouette Score
Cluster ID
Survival Group
Castration Status
Metastasis
Cluster
Cluster-specific Pathways
Cluster ID
Castration Status
Castrated
Intact
Metastasis
Localized
Metastatic
Pathway Enrichment
HALLMARK_INTERFERON_GAMMA_RESPONSE
KEGG_MATURITY_ONSET_DIABETES_OF_THE_YOUNG
REACTOME_INTERFERON_GAMMA_SIGNALING
REACTOME_REGULATION_OF_BETA_CELL_DEVELOPMENT
REACTOME_NUCLEAR_RECEPTOR_TRANSCRIPTION_PATHWAY
REACTOME_REGULATION_OF_GENE_EXPRESSION_IN_BETA_CELLS
HALLMARK_ALLOGRAFT_REJECTION
KEGG_HEMATOPOIETIC_CELL_LINEAGE
HALLMARK_PANCREAS_BETA_CELLS
HALLMARK_BILE_ACID_METABOLISM
HALLMARK_INTERFERON_ALPHA_RESPONSE
HALLMARK_ADIPOGENESIS
REACTOME_TRAF6_MEDIATED_IRF7_ACTIVATION
REACTOME_RIG_I_MDA5_MEDIATED_INDUCTION_OF_IFN_ALPHA_BETA_
KEGG_RIG_I_LIKE_RECEPTOR_SIGNALING_PATHWAY
KEGG_ADIPOCYTOKINE_SIGNALING_PATHWAY
HALLMARK_MYC_TARGETS_V1
HALLMARK_E2F_TARGETS
HALLMARK_G2M_CHECKPOINT
REACTOME_CELL_CYCLE
REACTOME_CELL_CYCLE_MITOTIC
HALLMARK_MYC_TARGETS_V2
REACTOME_S_PHASE
REACTOME_G1_S_TRANSITION
REACTOME_DNA_REPLICATION
REACTOME_METABOLISM_OF_RNA
REACTOME_MITOTIC_M_M_G1_PHASES
REACTOME_SYNTHESIS_OF_DNA
KEGG_DNA_REPLICATION
REACTOME_DNA_STRAND_ELONGATION
REACTOME_CHROMOSOME_MAINTENANCE
REACTOME_TELOMERE_MAINTENANCE

**Figure S4: Additional transcriptomic analyses of the GEMM cohort (related to Figure 3)**

**A.** Bioactivity score comparison using the GEMM interactome reported in Aytes et al 2014 (Old) the new one reported herein (New). The graph shows the probability density of the absolute value of the Normalized Enrichment Scores (NES) for the top 50 candidate MRs, across the 91 GEMM-DTs (GEMM-DT cohort). This shows that the New interactome is more effective at identifying proteins that are differentially active ($P < 10^{-16}$). **B,C.** Uniform manifold approximation and projection (UMAP) representation of the GEMM-DT cohort. **B.** Colors represent the distribution of the phenotypic variables shown in Figure 3 across all GEMM-DTs. **C.** Colors represent the distribution of protein activity for key markers, as identified in Figure 3, across all GEMM-DTs**.** **D.** Protein activity-based pathway analysis of the GEMM-DT cohort showing the most statistically significant pathways from the KEGG, REACTOME and Broad MSigDB Hallmarks of Cancer collections, as well as the relevant phenotypic variables discussed in Figure 3A.

**Figure S5**

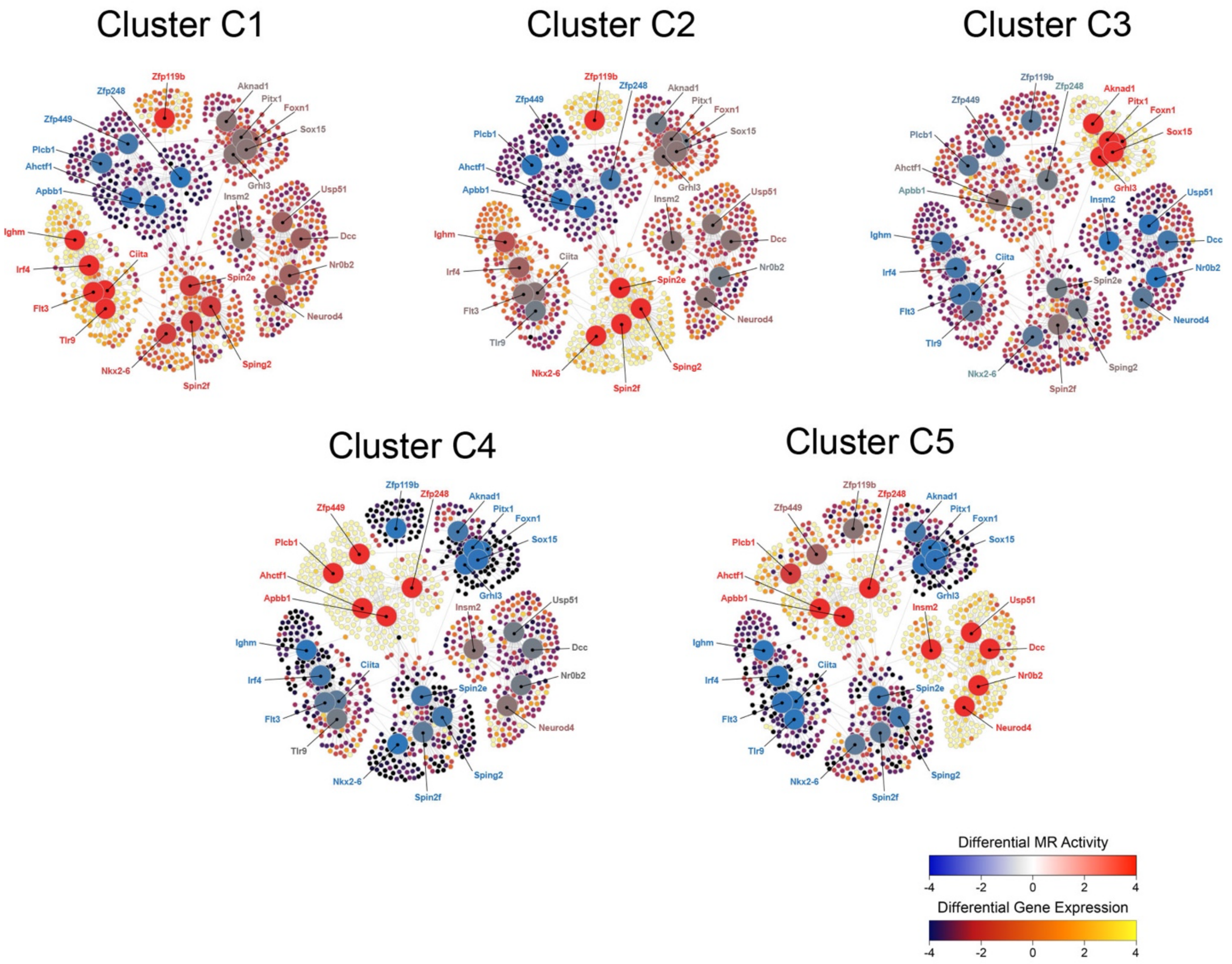


**Figure S5: Regulatory sub-networks of the GEMM clusters (related to Figure 3)**

High resolution view of the sub-networks shown in Figure 3B for each cluster ($C_1 - C_5$). See Figure 3B for details about representation and color scales.

**Figure S6**

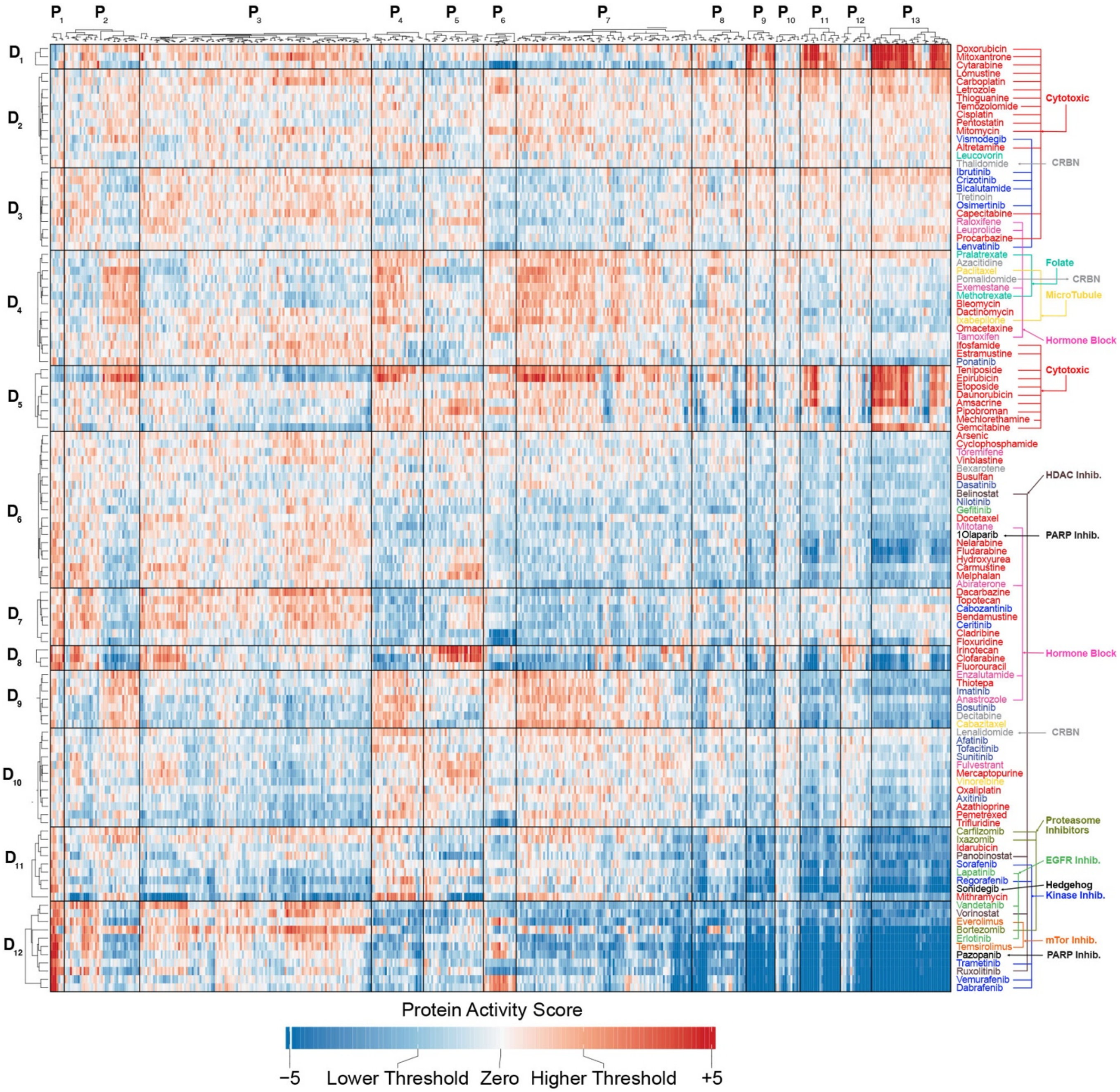


**Figure S6: Drug perturbation protein activity profiles from DU145 cells (related to Figures 5, 6, and 7)**

The heatmap shows the differential activity of protein (columns) in DU145 cells following perturbation with 117 FDA-approved drugs (column), using a blue (inactivated) to red (activated) color scale. For visualization purposes, only the 5% proteins with the greatest differential activity across all drug perturbations (high-variability proteins, HVP) are shown. Both proteins and drugs are clustered to show protein programs consistently activated or inactivated by specific drug subsets, see Methods. Drug families are indicated to the right of the plot showing both significant reproducibility across drugs targeting the same mechanism (e.g. EGFR or microtubule inhibitors) as well as the cell's ability to canalize the activity of drugs with different mechanism of action towards the same transcriptional cell state.

**Figure S7**

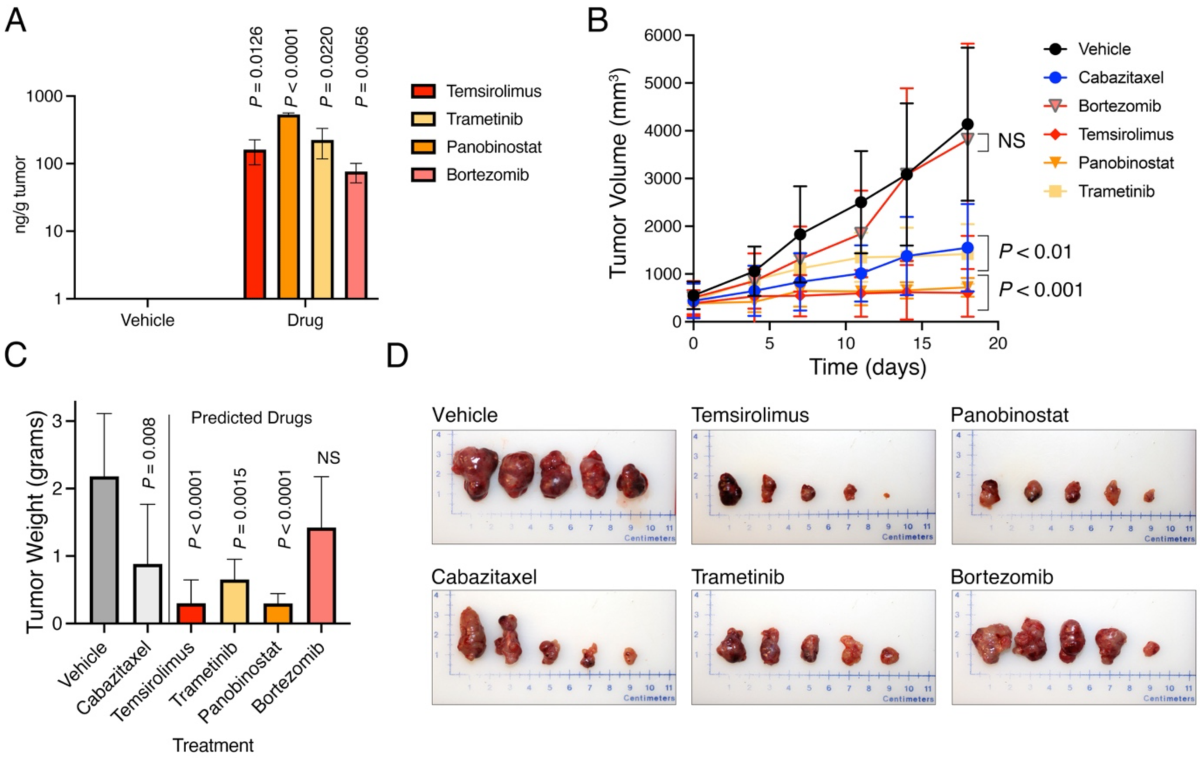


**Figure S7: Additional validation of drug candidates (related to Figures 6,7)**

**A.** Pharmacokinetic analysis: Intratumoral drug concentrations measured by UPLC-MS/MS, at the final timepoint of the experiment in the CMZ150 NPM allograft. **B-D.** Preclinical validation results: Validation of candidate drugs, *in vivo*, in the CMZ150 NPM allograft derived from a second cognate GEMM-DT (CMZ150). Allografts were grown subcutaneously in *nude* mouse hosts and treated with predicted drugs, negative control (cabazitaxel) and vehicle control, for the times indicated. **B.** Summary of changes in tumor volume over the treatment period. **C.** Summary of tumor weights following sacrifice. *P*-value analysis and animal per drug arm are as reported in Figures 6, 7. **D.** Representative images of final tumor sizes.
